# Multi-omics Topic Modeling for Breast Cancer Classification

**DOI:** 10.1101/2021.12.22.473851

**Authors:** Filippo Valle, Matteo Osella, Michele Caselle

**Affiliations:** Physics Department, University of Turin and INFN, via P Giuria 1, 10125 Turin, Italy; (M.O.); (M.C.)

**Keywords:** miRNAs, miRNA expression regulation, topic modeling, stochastic block modeling, multi-omics, chr14q32

## Abstract

The integration of transcriptional data with other layers of information, such as the post-transcriptional regulation mediated by microRNAs, can be crucial to identify the driver genes and the subtypes of complex and heterogeneous diseases such as cancer. This paper presents an approach based on topic modeling to accomplish this integration task. More specifically, we show how an algorithm based on a hierarchical version of stochastic block modeling can be naturally extended to integrate any combination of ‘omics data. We test this approach on breast cancer samples from the TCGA database, integrating data on messenger RNA, microRNAs and copy number variations. We show that the inclusion of the microRNA layer significantly improves the accuracy of subtype classification. Moreover, some of the hidden structures or “topics” that the algorithm extracts actually correspond to genes and microRNAs involved in breast cancer development, and are associated to the survival probability.

**Simple Summary:** Topic models are algorithms introduced for discovering hidden topics or latent variables in large unstructured text corpora. Leveraging on analogies between texts and gene expression profiles, these algorithms can be used to find structures in expression data. This work presents an application of topic modeling techniques to the identification of breast cancer subtypes. In particular, we extended a specific class of topic models to allow a multi-omics approach. As an illustrative example, considering both messenger RNA and microRNA expression levels, we were able to clearly distinguish healthy from tumor samples, as well as the different breast cancer subtypes. The integration of different layers of information is crucial for the observed classification accuracy. Our approach naturally provides the genes and the microRNAs associated to the specific topics that are used for the sample organization. We show that indeed these topics often contain genes involved in breast cancer development, and are associated to different survival probabilities.

## 1. Introduction

A crucial problem in modern computational biology is the integration of different sources of information in the framework of the so-called “precision medicine” [1]. Thanks to the impressive improvement of experimental techniques and the creation of dedicated databases, plenty of different ‘omics data sets are available. However, these data sets are difficult to integrate in a coherent picture. They are typically noisy and sparse, they can strongly depend on experimental and processing choices and biases, such as normalization or imputation techniques, and can present different constraints for example due to (often unknown) specific regulatory interactions. At the same time, only by combining different layers of information we hope to understand complex pathologies like cancer and thus optimize the therapeutic protocols. In fact, a major goal would be to be able to identify as soon as possible the particular cancer subtype of a given patient, find the corresponding drivers and altered pathways, and thus, possibly, fine tune the therapy. A fundamental preliminary step is the development of algorithms able to identify and extract the relevant structure and organization of tumor samples using the different available layers of molecular information.

In particular, topic modeling has been recently proposed as a computational technique to identify hidden structures in gene expression data [2,3]. Topic models are a set of algorithms originally developed to extract latent variables from texts corpora [4–6]. The most popular of these algorithms is the so-called Latent Dirichlet Allocation [5] (LDA) which has been successfully applied not only in texts analysis, but also in other contexts such as bioinformatics [7].

LDA is based on the assumption of a Dirichlet prior for the latent variables. This choice simplifies the statistical inference problem making the algorithm highly efficient. However, many complex systems in which LDA is applied, including expression data, are characterized by the emergence of power-law distributions which are very far from the Dirichlet assumption [8–11]. Moreover, the optimal number of topics must be identified by the user in the standard LDA formulation [5].

To overcome these problems, a new class of algorithms based on hierarchical Stochastic Block Modeling (hSBM) was recently proposed [10]. These algorithms are based on the formal equivalence between the topic identification problem and the community detection problem in bipartite networks [12–14], where well developed techniques based on stochastic block modeling [15] can be applied without the need of a Dirichlet prior.

We recently performed a comparative study [3] of different topic modeling algorithms on the task of identifying cancer subtypes from breast and lung cancer gene expression datasets from The Cancer Genome Atlas (TCGA) [16,17]. We found that hSBM typically outperforms other algorithms in the clustering task. Importantly, this algorithm presents the additional advantages of naturally selecting the number of clusters and of providing the genes significantly associated with the latent structure on which the classification is based. We were able to show that the established cancer subtype organization for both breast and lung cancer was well reconstructed by the latent topic structure inferred by hSBM and that the topic content itself was very informative. In fact, topics associated with specific cancer subtypes were enriched in genes known to play a role in the corresponding disease, and were related to the survival probability of patients.

This paper extends our previous study by integrating in the hSBM framework multiple layers of information. While the integration of additional biological information should generally improve the accuracy of the statistical inference, it is important to stress that this is not always trivially true. Highly noisy or irrelevant data layers could interfere with the task. We will show an empirical example of such a negative interference. Therefore, the addition of new layers should be driven by a clear biological motivation.

We will focus on the illustrative case of breast cancer, which is the most commonly diagnosed cancer type and the leading cause of cancer death in women worldwide [18], with three main goals:

- First, we will show how different layers of biological information can be efficiently integrated in the hSBM framework. We release the python package *nSBM*, inherited from hSBM [10], which is ready to install, easily executable, and can be used to infer the topic structure starting from different layers and type of biological data.
- Second, focusing on breast cancer, we will show that the combination of microRNA and protein-coding expression levels greatly improves the algorithm ability to identify cancer subtypes. These findings further confirm the important role previously recognized in several studies that miRNAs play in cancer development [19,20].
- Third, we use the inferred topic structure to select few genes, miRNAs and chromosomal duplications which seem to have a prognostic role in breast cancer and thus could be introduced as additional signatures of specific breast cancer subtypes. The extension of subtype signatures can help clinicians to fine tune diagnostic protocols in the framework of a precision medicine approach to cancer [1].

## 2. Results

### 2.1. nSBM: a multi branch topic modeling algorithm

Many real-word networks are accompanied by annotations or metadata describing different node properties. For example in social networks, information about age, gender or ethnicity can be associated to the nodes, or the data capacity can be associated to the nodes of the Internet network [21]. In a similar way, different ‘omics can provide additional information to biological networks. These metadata can improve the performance of community detection algorithms by providing additional levels of node correlations that are not accessible only using a single data source [22–24]. Given the relation between community detection and topic modeling [10], a similar improvement is expected also in the detection of latent variables using topic modeling analysis on multi-omics data sets. Our first goal is thus to extend the topic modeling approach to multi-omics data, and to test its performances in a concrete biological problem.

The extension of a network-based topic modeling algorithm to multi-partite networks was recently proposed in the classic context of texts analysis by [23], and we apply here a similar approach to biological data. In this case networks are generic n-partite networks that contain nodes of n types: sample nodes (i.e. patients), and (n-1) sets of nodes (e.g. protein-coding mRNA levels, microRNAs, etc.) which represent different features associated to the sample nodes.

The topology of the n-partite network is star-like with a center containing the sample nodes and *n* – 1 branches (Figure 1b). Each node in a branch can be connected with all the sample nodes, but no connection exists between nodes within a branch nor between nodes in different branches. This is the natural generalization of the standard bipartite network shown in Figure 1a. In the biological example that will be addressed in the following only two branches are present: protein-coding genes and microRNAs. However, the presented scheme is general and can be easily extended to several branches at the expense of computational speed. We will discuss the addition of a third sample feature capturing the gene Copy Number Variation (CNV).

**Figure 1.**
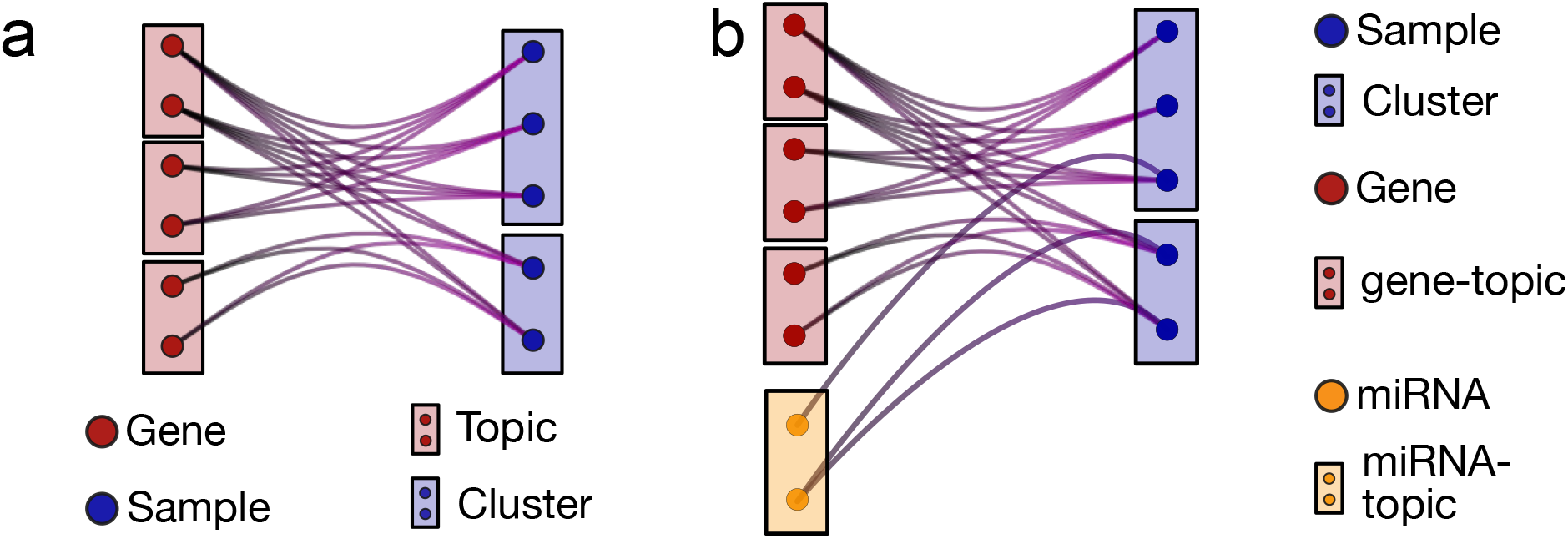
Cartoon of multi-partite networks with samples, protein-coding genes and micro-RNAs. (**a**) A bi-partite network with a layer of protein-coding genes and a layer of samples. A gene is connected to a sample if it is expressed in that sample and the link weight is proportional to the expression level. (**b**) A tri-partite network obtained by adding the miRNA expression layer. The topic model algorithm essentially outputs a block or topic structure in each layer.

We shall denote in the following as “links” the connections between the branch nodes and the sample nodes. Each link is characterized by a weight. The weights can have a different nature depending on the branch. For instance, weights on links connecting the gene branch with the samples encode the expression level (here in FPKM units), and analogously the links connecting to miRNAs report the miRNA expression level. When we add a layer with the CNV information, the links are weighted with the number of copies of the gene in the connected sample. The algorithm interpret the weight *w_ij_* between node *i* and node *j* as a collection of *w_ij_* independent edges. We will use the term “edge” for this elementary unit of link weights.

Once the multi-partite network is defined, the statistical inference procedure leading to the topic structure is a straightforward extension of the procedure developed for the hierarchical Stochastic Block Model (hSBM) [10], which we already applied in its bipartite form to expression data [25]. hSBM is a generative model that basically searches the parameters (*θ*) that maximise the probability that the model describes the data 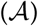

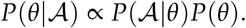

The model uses a generative process to build a network given a set of parameters *θ*. Using a Markov Chain MonteCarlo algorithm these parameters are optimised in a unsupervised way and the optimisation continues until the generated model approximates well the data 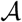. (see [10] and references therein for more details).

The output of the algorithm is a partitioning of nodes or a set of “blocks” of nodes associated to probability distributions. The samples are partitioned into “clusters”, while the blocks of nodes in the branches are essentially the “topics”. Since we are considering several branches, we will have topics of different types, such as gene-topics on the gene expression branch, miRNA-topics on the miRNA branch, CNV-topics on the CNV branch and so on. We will consider clusters and topics as “hard” blocks (i.e., each sample/gene/miRNA belongs to only one block) and distinct (there are no blocks containing different kind of nodes). However, given its probabilistic nature, the algorithm can be naturally extended to fuzzy clusters.

There are several features that distinguish hSBM, and its nSBM extension introduced here, from other clustering or topic modeling algorithms such as LDA.

- *Lack of a parametric prior*. Thanks to the network-based approach and to the particular way links are used to update the block structure, this class of algorithms does not require a specific parametric assumption for the prior probability distribution of the latent variables. This is a major difference with respect to LDA, and makes this class of algorithms particularly suited for biological systems in which long-tail distributions and hierarchical structures are ubiquitous (see the discussion on this point and the comparison with LDA in [3]).
- *Probability distributions over latent variables of different types*. The output of the algorithm is not deterministic but is instead a set probabilities that associate a sample with latent variables of different types *P*(gene-topic|sample), *P*(miRNA-topic|sample), and that associate different features to topics, such as *P*(gene|gene-topic) and *P*(miRNA|miRNA-topic). *P*(gene-topic|sample) and *P*(miRNA-topic|sample) represent the contribution of each miRNA- or gene-topic to each sample. On the other hand, *P*(gene|gene-topic) and *P*(miRNA|miRNA-topic) quantify how much each gene or miRNA contributes to a specific topic. As we will show in the following, these probability distributions capture relevant properties of the biological system.
- *Hierarchical topic structure*. Blocks and the probability distributions described above are available at different layers of resolution, from few large sets (clusters/gene-topics/miRNA-topics) at low resolution, to many small sets at a higher resolution. The specific number of layers and their block composition are found by the algorithm optimization process, and are not given as input. Therefore, the data sets can be organized in different ways depending on the resolution of interest. Note that not all possible resolutions are trivially present, as in standard hierarchical clustering.
- *Concurrent and separate topic organization of the different network layers*. Different ‘omics have typically different normalisation and the numbers associated to different molecular features have often a very different meaning. A major advantage of nSBM with respect to other algorithms [26,27] is that each layer is independently contributing to the optimization process and a topic organization is given for each layer. Therefore, there is no need to re-weight the different layers to balance their contributions since they are kept separate while concurrently contributing to the sample clustering. This makes the model suitable to be applied not only to genomics data, as we will discuss in this paper, but, ideally, to any combination and number of different concurrent ‘omics.

### 2.2. Subtype classification of breast cancer samples

The benchmark task we now focus on to test the performance of nSBM is its ability to cluster breast cancer samples according to their subtype annotation. This is an important task for its clinical relevance, but also because the breast cancer subtype could be dependent on a complex combination of factors, including gene and miRNA expression profiles, and thus the classification could be a good test for nSBM.

Breast cancer is indeed a heterogeneous disease, with wide variation in tumour morphology, molecular characteristics, and clinical response [18,28–30]. Notwithstanding this variability, it is one of the few tumors for which there is a widely accepted subtype classification [28,31].

Breast cancer samples are usually divided in 5 different subtypes: *Luminal A, Luminal B, Triple-Negative/Basal, HER2* and *Normal-like*. For our tests, we used as benchmark the TCGABiolinks annotations [32,33], as discussed in the Methods section. These annotations are the result of a rather complex process. On the clinical side the classification is based on the levels of a few proteins whose presence in the biopsy is usually detected using immunohistochemistry (IHC) assays. In particular, these proteins are two hormone-receptors (estrogen-receptor (ER) and progesterone-receptor (PR)), the Human Epidermal growth factor Receptor 2 (HER2), and Ki-67, which is a nuclear antigen typically expressed by proliferating cells and thus used as an indicator of cancer cells growth. On the gene expression side, the same subtypes can be identified by looking at the expression levels of a set of genes included in the so called “Prediction Analysis of Microarray (PAM)50” [34]. The agreement between PAM50 results and IHC-based subtyping is in general reasonably good, but far from being perfect. Indeed, the classification task is made particularly difficult by the heterogeneity of cancer tissues (biopsies may contain relevant portions of healthy tissue) and by the intrinsic variability of gene expression patterns in cancer cell lines.

We recently demonstrated that topic modeling based algorithms can achieve satisfactory performances in this classification task by looking at gene expression profiles [3] (and not only of the PAM50 genes), and not relying on the known IHC markers. The advantage of this approach is that it avoids problems and ambiguities in the classification due to the stochastic fluctuations of the IHC markers or due to the different inference strategies adopted by PAM50 classifiers (see for instance [35] for a recent comparison of the performances of different classifiers in a set of breast cancer classifications tasks).

Following this line, one of the goals of our study is to evaluate if the integration of miRNA expression levels (and possibly of other layers of information) can further improve the hSBM results presented in ref. [3].

### 2.3. Integrating microRNA expression profiles in a topic modeling analysis

It is, by now, well established that miRNAs play an important role in several human diseases and in particular in cancer. Accordingly, miRNAs have been proposed as diagnostic biomarkers of human cancers [20,36,37]. This is particularly true for breast cancer, for which several studies highlighted the prognostic role of miRNAs [38].

Following this line of evidence, we integrated miRNA expression levels with protein-coding mRNA levels using a *n* = 3 version of nSBM (which in the following we shall denote as triSBM). In this case, the analysis output, besides the clusters of samples and the topics of genes, will also contain a collection of miRNA-topics.

#### 2.3.1. Including miRNAs in the topic modeling analysis modifies both the sample clusters and the gene-topics

We first tested if the integration of miRNAs have an effect on the partition of samples in clusters and in the topic organization in the gene branch.

Figure 2 reports the Adjusted Mutual Information (AMI) between the partition obtained with a standard hSBM and with triSBM, varying the hierarchy level (*l*0, *l*1…), being *l*0 the finer layer (the one with smaller sets). We used the AMI to score the overlaps of partitions, since it measures the mutual information between partitions compared to the one obtained by two random partitions. Figure 2a shows that there is a substantial disagreement between the clusters of samples in the two outputs. Similarly, Figure 2b indicates that the same is true for the topics on the protein-coding gene side. The overlap between the partitions obtained by hSBM and triSBM is negligible.

**Figure 2.**
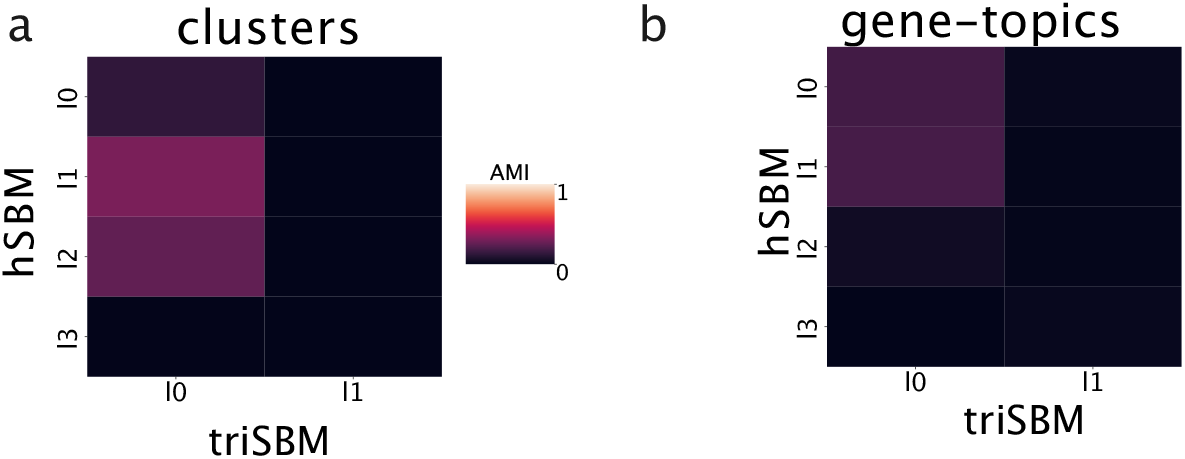
Adding miRNA leads to new topics. The Adjusted Mutual Information between the outputs of triSBM and hSBM (i.e. with and without miRNA). The partition obtained in output are different for any combination of layers.

Therefore, the addition of the miRNA branch can radically affect the inferred topic structure and the clustering of samples.

#### 2.3.2. The inclusion of miRNAs in the topic modeling analysis leads to a better separation of healthy and tumor tissues

We first test the ability of the algorithm in recognizing healthy from cancer samples. The hSBM algorithm showed good performances on this task by considering only gene expression data [3], as summarized in Figure 3b. We now test triSBM, in which gene expression levels are considered jointly with miRNA levels in the same set of TCGA samples. The detailed procedure and the algorithm output at different hierarchical levels are described in the Methods section. However, we basically found a significant improvement in the performance of the algorithm. In fact, Figure 3a clearly shows that normal samples are collected in a single cluster by triSBM, while the separation is less neat in absence of information on miRNA expression (Figure 3b)

**Figure 3.**
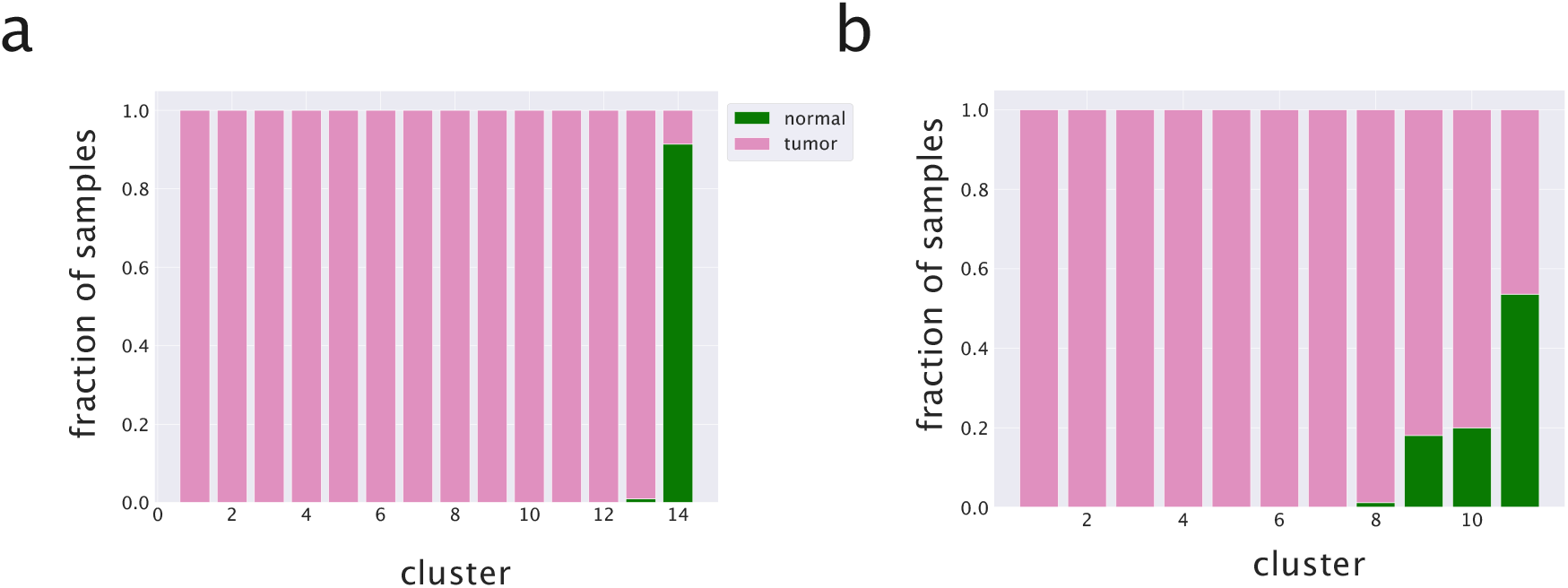
Clustering of breast samples with and without the miRNA branch. We compare normal and solid tumor tissues from TCGA using (**a**) triSBM and (**b**) hSBM at a similar resolution level.

The two model settings (hSBM and triSBM) are compared quantitatively in Figure 4 using the Normalized Mutual Information (NMI) as a score [39,40]. The NMI score is explained in detail in the Methods section.

**Figure 4.**
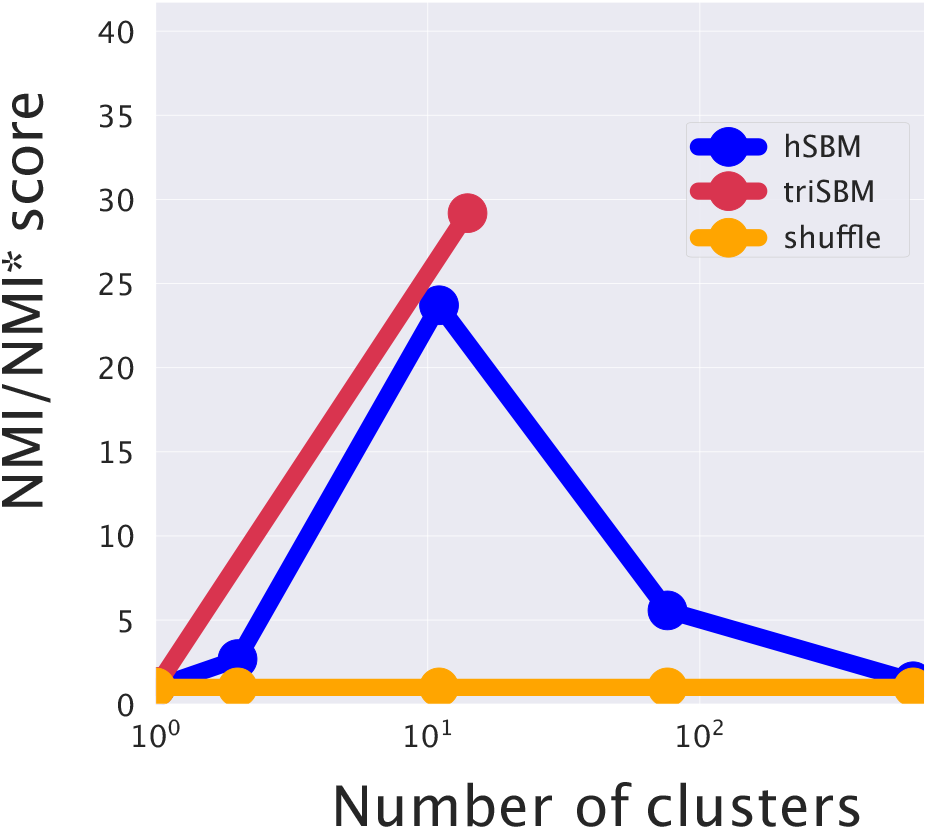
The increase in performance in separating tumor and normal samples by the addition of the miRNA layer. The NMI is evaluated at different resolution levels (number of clusters) using (triSBM) or not using (hSBM) the information of miRNA expression. The normal/tumor annotation from TCGA is used as ground truth.

#### 2.3.3. Including miRNAs in the topic modeling analysis improves the identification of cancer subtypes

As a second benchmark we tested the ability of triSBM to identify breast cancer subtypes. Again triSBM and hSBM are compared and the results are reported in Fig 5. Also in this case, the inclusion of miRNA levels improves the algorithm ability to group samples belonging to the same cancer subtype. The improvement is quantified by the NMI score reported in Fig. 5a. Fig. 5b and 5c show that the improvement is mainly due to a better performance of triSBM in distinguishing LuminalA from LuminalB samples. This was indeed the critical obstacle limiting the performances of hSBM in our previous study [3], suggesting that the distinction of these subtypes crucially depend on miRNA expression levels.

**Figure 5.**
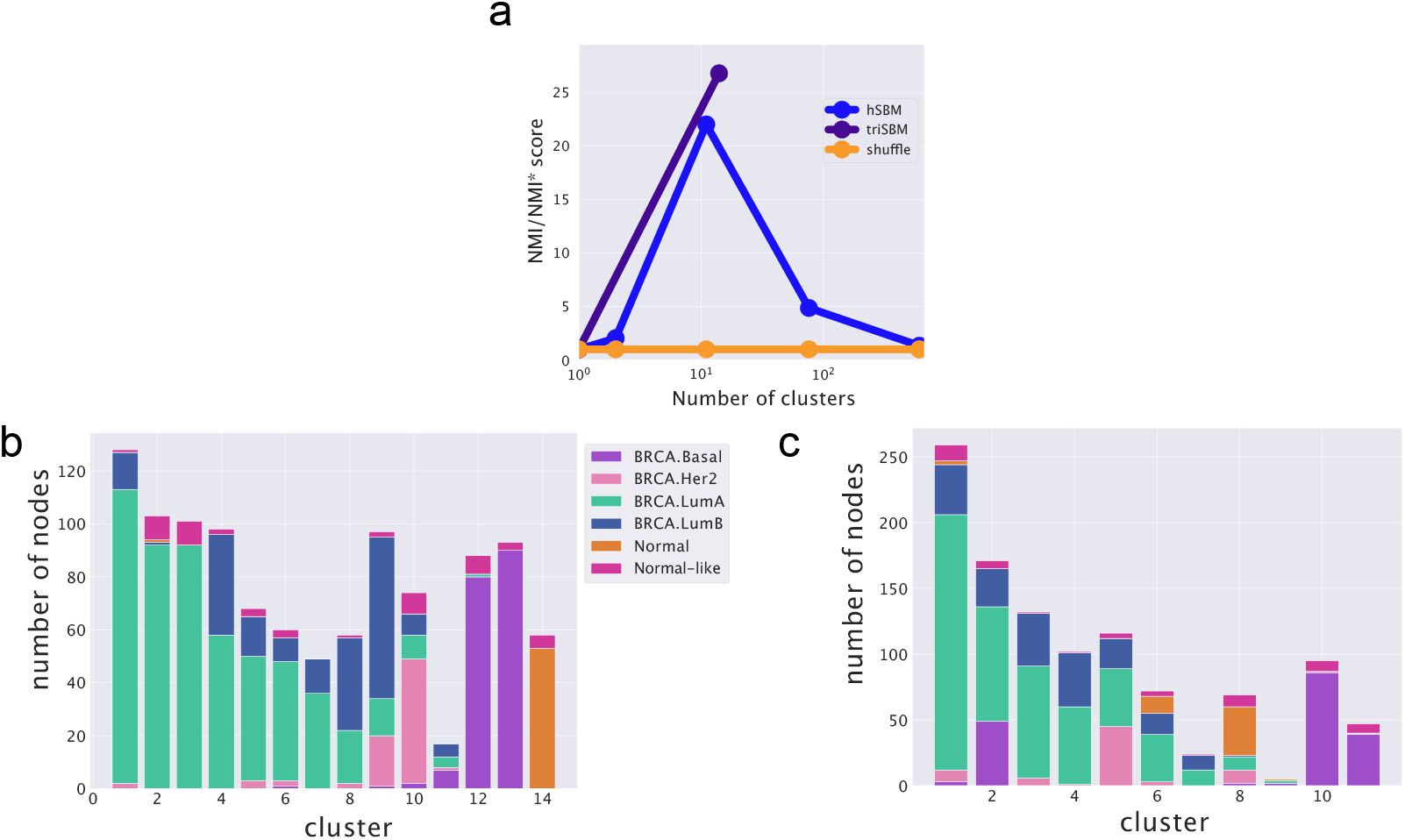
Scores and partitions based on *Subtype_Selected* annotations from [32,33]. (**a**) Scores for both (triSBM and hSBM) setting for the subtype classification problem. In (**b**) the miRNA were introduced. We compared the two settings choosing the layers with a compatible number of clusters. In (**c**) the clusters from a simple bi-partitie setting. They are almost similar, in (**c**) Luminal B is identified better. We define Normal the *Solid Tissue Normal* from TCGA, whilst Normal-Like are the *Primary Tumors* annotated BRCA.Normal from [32].

We used the *Subtype Selected* labels provided by TCGABiolinks [32,33] as the ground-truth annotation of subtypes. Note however that this labelling has a less solid basis with respect to the clear healthy/cancer distinction since the subtypes may not be so clearly defined and can be easily mis-classified because of the high tumor heterogeneity.

Note that the standard characterization of breast subtypes relies on the expression level of only few markers. We did not explicitly selected these markers in our gene selection process and thus, as previously discussed [3], the emergent sample organization is the result of the global pattern of gene and miRNA expression levels. Therefore, the significant overlap with the standard subtype annotation is highly non trivial, and the discrepancy does not have to automatically interpreted as a failure since the standard annotation could be limited.

Given these positive results, we will explore in the following sections the biological information contained in the latent variables inferred by the algorithm, and test their possible prognostic role.

#### 2.3.4. Check the robustness of the model with an independent labeling

We compared the blocks we obtained in output with the annotations of TCGA sample in [41]. First of all we measured the Adjusted Mutual Information (AMI) between these labels and the Subtype_Selected ones discussed above (AMI is a score between 0 and 1, which measures the mutual information between two annotations compared to the one obtained comparing two random annotations). We found a value of ~0.37, which shows that the two labelling are not trivially the same and thus represent a reliable test of our clusters.

We measured the Normalised Mutual Information score of both the bi-partite (hSBM) model and the model which integrates miRNA (triSBM). Results are reported in Supplementary Figure S1. Looking at the figure we see that our clusters show a significant agreement (high values of NMI/NMI*) also with this independent classification and, above all, that the agreement improves including miRNAs.

The overlap between our cluster partition and two independent non overlapping labels can be explained by the fact that our partition groups samples at the intersection between the two labelling system.

#### 2.3.5. Validation on an independent source of data: METABRIC

We applied the same pipeline applied on TCGA to the METABRIC [42] dataset and we measured the agreement between our partition on this data and the labels provided by [41]. We confirmed the results obtained on TCGA: the triSBM model has a better agreement (NMI score is reported in Supplementary Figure S2) with the labels assumed as ground truth with respect to the model without miRNA (hSBM).

### 2.4. triSBM topics can be used to obtain subtype-specific information

A major advantage of a topic modeling approach to multi-omics data is that we can use the information stored in the probability distributions *P*(topic|sample) to obtain sub-type specific signatures. Following the analysis of [3] we constructed from these probabilities a set of “centered” distributions 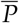(topic|subtype) (see the definition in the Equation 5 of the Methods section) which allow us to identify subtype specific topics (i.e. topics which are particularly enriched in the samples belonging to a particular subtype) that are thus candidates to play a role in driving the specific features of that subtype.

These topics are nothing but lists of genes and can be investigated using a standard enrichment analysis. The results shown in this paper were computed using the Gene Set Enrichment Analysis GSEA [43] tool. In particular, we concentrated on the keywords extracted from [44], [45] and [46].

We discuss the results of this analysis in the following two subsections.

#### 2.4.1. Analysis of subtype specific topics of genes

We report in Figure 6 a few examples of 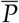(gene-topic|subtype) distributions for a few selected topics and in Table 1 the results of the corresponding enrichment analysis.

**Figure 6.**
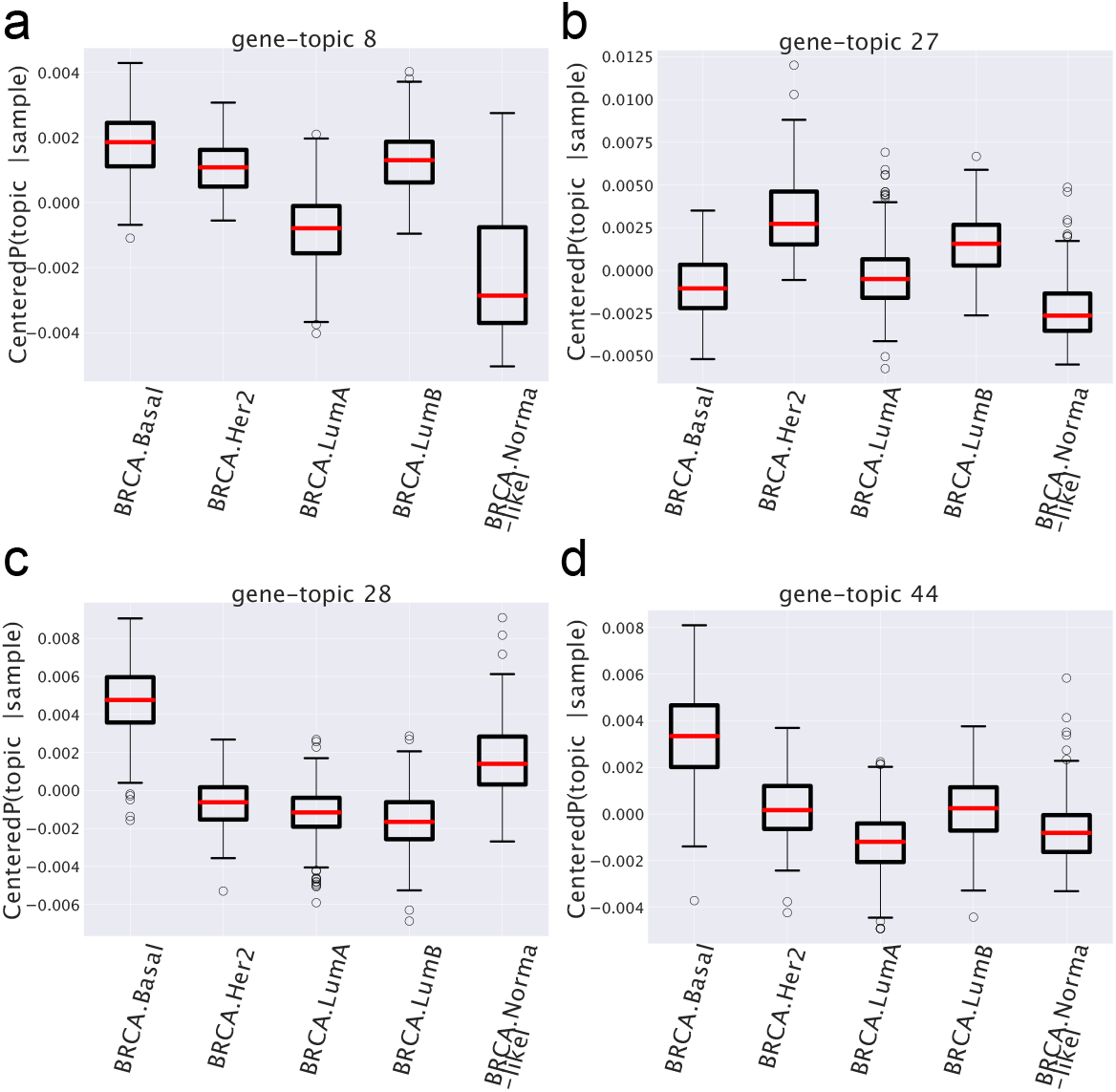
Box plots of the centered *P* (gene-topic|sample) for different gene-topics. Samples belonging to each subtype may be over or under expressed in different topics.

**Table 1.** GSEA FDR enrichment P-values on the gene-topics. For each gene-topic only the terms with the strongest enrichment are reported. In brackets the number of genes in each set (topic). Lists are available at https://github.com/BioPhys-Turin/keywordTCGA/blob/main/brca/trisbm/trisbm_level_0_topics.csv.

| Term | False Discovery Rate |
| --- | --- |
| gene-topic 6 (55) |  |
| SMID_BREAST_CANCER_BASAL_DN | $8.14 * 10^{-22}$ |
| FARMER_BREAST_CANCER_APOCRINE_VS_LUMINAL | $3.67 * 10^{-7}$ |
| gene-topic 8 (19) |  |
| MODULE_54 (cell cycle) | $2.31 * 10^{-20}$ |
| gene-topic 12 (13) |  |
| MODULE_1 (ovary genes) | $1.52 * 10^{-7}$ |
| SMID_BREAST_CANCER_BASAL_DN | $1.53 * 10^{-7}$ |
| gene-topic 15 (26) |  |
| HALLMARK_EPITHELIAL_MESENCHYMAL_TRANSITION | $7.93 * 10^{-15}$ |
| gene-topic 25 (40) |  |
| CHARAFE_BREAST_CANCER_LUMINAL_VS_MESENCHYMAL_UP | $4.59 * 10^{-15}$ |
| SMID_BREAST_CANCER_BASAL_DN | $3.98 * 10^{-14}$ |
| VANTVEER_BREAST_CANCER_ESR1_UP | $7.91 * 10^{-5}$ |
| gene-topic 27 (44) |  |
| SMID_BREAST_CANCER_ERBB2_UP | $1.73 * 10^{-7}$ |
| gene-topic 28 (53) |  |
| SMID_BREAST_CANCER_BASAL_UP | $8.11 * 10^{-23}$ |
| gene-topic 37 (54) |  |
| FAN_OVARY_CL13_MONOCYTE_MACROPHAGE | $1.36 * 10^{-14}$ |
| VANTVEER_BREAST_CANCER_ESR1_DN | $7.63 * 10^{-11}$ |
| gene-topic 44 (37) |  |
| SMID_BREAST_CANCER_BASAL_UP | $3.22 * 10^{-13}$ |
| gene-topic 53 (39) |  |
| SMID_BREAST_CANCER_BASAL_DN | $4.46 * 10^{-11}$ |
| gene-topic 55 (58) |  |
| SMID_BREAST_CANCER_BASAL_DN | $4.96 * 10^{-33}$ |
| FARMER_BREAST_CANCER_BASAL_VS_LUMINAL | $1.21 * 10^{-14}$ |
| VANTVEER_BREAST_CANCER_ESR1_UP | $2.91 * 10^{-14}$ |
| SMID_BREAST_CANCER_LUMINAL_B_UP | $7.33 * 10^{-12}$ |
| gene-topic 68 (54) |  |
| SMID_BREAST_CANCER_BASAL_UP | $6.19 * 10^{-21}$ |
| CHARAFE_BREAST_CANCER_LUMINAL_VS_BASAL_DN | $1.32 * 10^{-14}$ |
| SMID_BREAST_CANCER_LUMINAL_B_DN | $1.46 * 10^{-14}$ |
| CHARAFE_BREAST_CANCER_LUMINAL_VS_MESENCHYMAL_DN | $3.15 * 10^{-10}$ |

Looking at the figures and at the table we see a few interesting patterns.

- There are topics, like for instance topic 8 in Figure 6 which show a similar behaviour in all cancer subtypes and a different one (in the case of topic 8, it is depleted) in the normal tissues. These are the topics which allowed the algorithm to distinguish so accurately normal from cancer samples. In the case at hand the functional analysis allows to easily understand the reason of this different behaviour: the genes contained in topic 8 are strongly enriched in cell cycle keywords, which is likely to be associated to the proliferating nature of tumour tissues.
- Another interesting pattern is well exemplified by topics 27,28 and 44 in Figure 6. These are topics which are over-represented only in one particular subtype (in the example topic 28 and 44 in the Basal subtype and topic 27 in the HER2 one) and can thus be used as signatures of these subtypes. This is in nice agreement with the finding of the gene enrichment analysis which for topics 28 and 44 gives a strong enrichment for the keyword SMID_BREAST_CANCER_BASAL_UP which is known to be associated to the Basal subtype [44], while topic 27 is enriched in the keyword: SMID_BREAST_CANCER_ERBB2_UP which is in fact associated to the HER2 subtype [44]. These topics are the latent variables which allow the algorithm to distinguish among different subtypes.

#### 2.4.2. Analysis of subtype specific topics of miRNAs

While the above results were similar to the ones already discussed in [3], the novelty of the present analysis is that we can perform a similar study also on the miRNA side. As we shall see this allows to have a new independent insight on the problem.

We report four instances of the 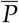(miRNA-topic|subtype) probability distributions in Figure 8 and the corresponding enrichment analysis in Table 2. They are, somehow, paradigmatic examples of the type of information which one can obtain from this type of analysis.

- The first one (named miRNA-topic 7 in our output, see: https://github.com/BioPhys-Turin/keywordTCGA/blob/main/brca/trisbm/trisbm_level_0_topics.csv) is the typical example of a topic which shows no particular preference for a cancer subtype (see Figure 6) but shows a strong enrichment for a particular chromosmal locus: chr14q32 (see Table 2). This enrichment is due to the fact that most of the miRNAs of the topic are indeed contained in this locus. Moreover, we see, looking at Figure 7, that these miRNAs are exactly those with the highest probability to belong to the topic. This strongly suggests that a somatic alteration (duplication or deletion) at this locus could be associated to the onset of cancer and could thus be used as a marker. Indeed, this locus is known to be associated to breast cancer [47]. Accordingly, if we perform a survival analysis, between patients with this topic upregulated and patients with the topic downregulated (see next subsection) we find a remarkable increase in the survival probability of patients with the topic downregulated.

**Figure 7.**
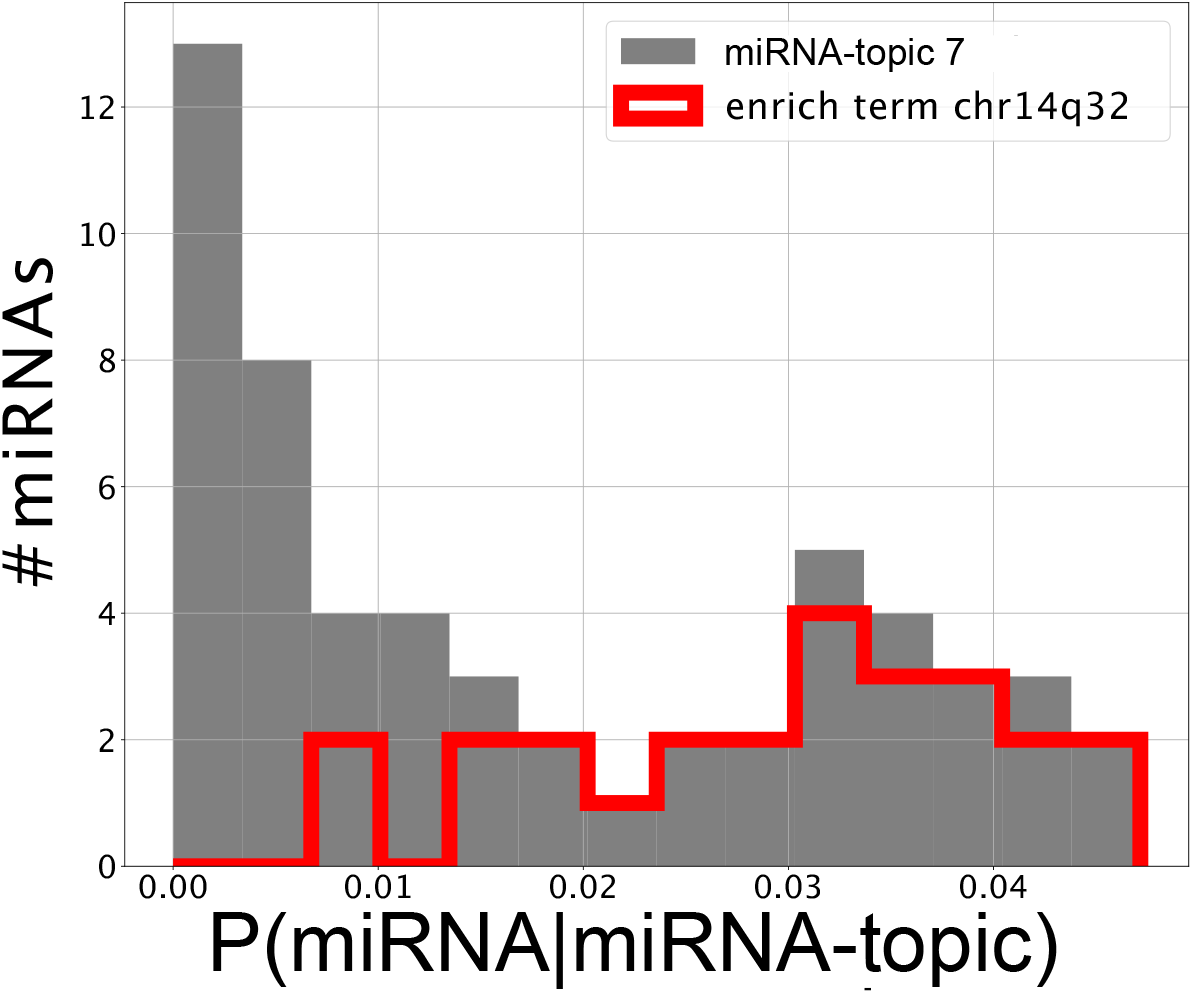
Genes that are annotated in the Gene Set Enrichment Analysis terms contribute more than the average to the topic. Contribution of miRNAs to miRNA-topic 7. miRNAs that belong to the ontology specific of locus *c14q32* are highlighted and have high *P*(miRNAjmiRNAs’ topic).

**Figure 8.**
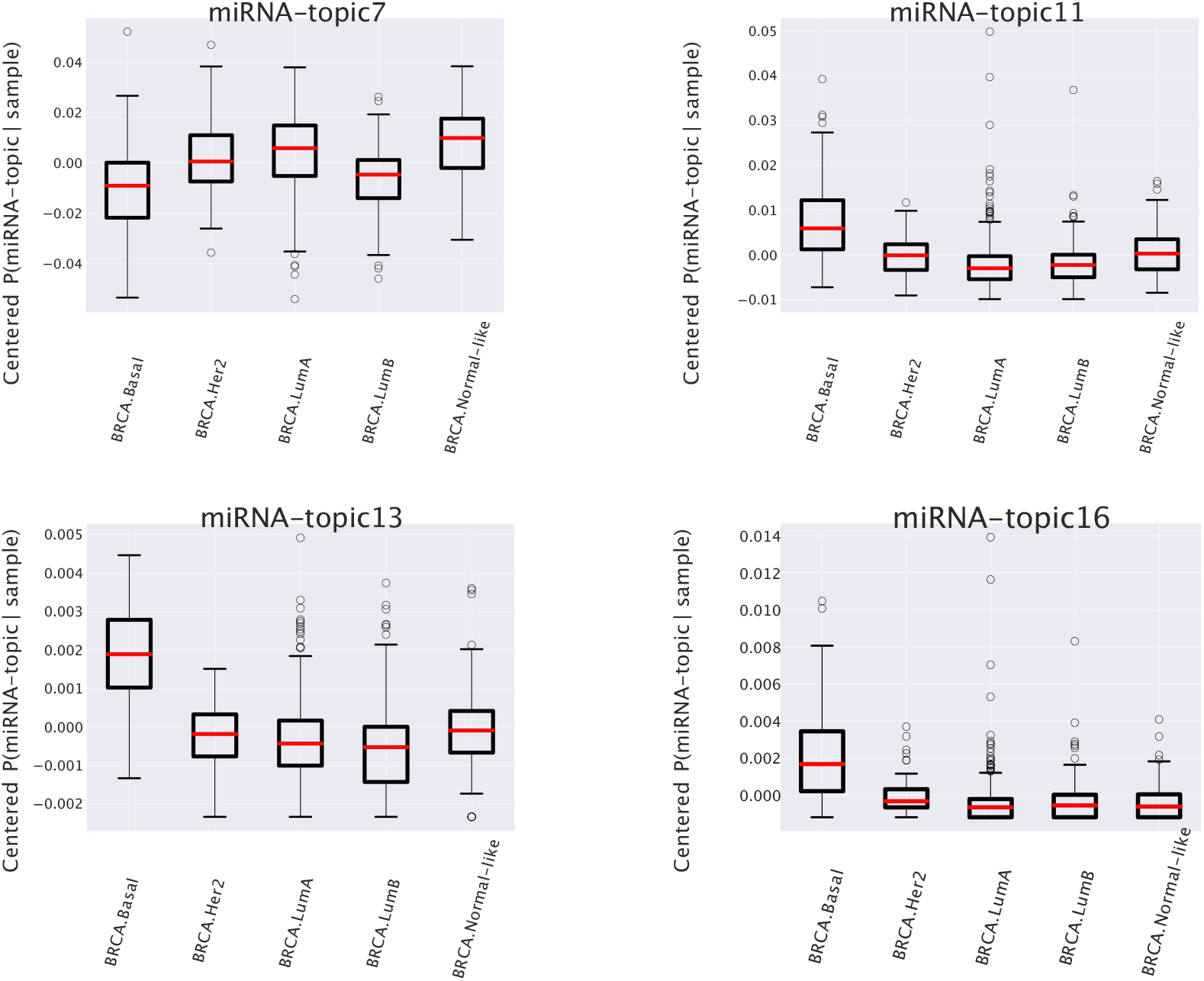
Box plots of the centered *P*(miRNA-topic|sample). This plot shows that the differences of topic expression in each subtype may be different. Some miRNA-topics are more abundant in samples known to be Basal Subtype.

**Table 2.** GSEA results on the miRNA-topics. We selected and reported the ones with strongest enrichment. Lists are available at https://github.com/BioPhys-Turin/keywordTCGA/blob/main/brca/trisbm/trisbm_level_1_metadata.csv.

| Term | False Discovery Rate |
| --- | --- |
| miRNA-topic 7 (57) |  |
| chr14q32 | $1.81 * 10^{-29}$ |
| WP_ALZHEIMERS_DISEASE | $2.88 * 10^{-4}$ |
| miRNA-topic 11 (60) |  |
| chr19q13 | $1.57 * 10^{-6}$ |

However this is not the end of the story. Looking at Table 2 we see that the same topic is also enriched in keywords associated to Alzheimer desease. Indeed it is known that there is a sort of inverse co-morbidity [48] between a few types of cancer (in particular lung [49] and breast [50]) and the Alzheimer desease. This association is confirmed and supported by our analysis, which also suggests that it could be mediated exactly by the microRNAs contained in miRNA-topic 7. Indeed, some of the miRNAs contained in the topic, like mir-34c, are known oncosuppressors of breast cancer [51,52] and, at the same time, are recognized markers of the Alzheimer desease [53,54]. The most important of these is the above mentioned mir-34c, which is in fact, strongly associated to miRNA-topic 7, being the only miRNA in the topic with *P*(miRNA|miRNA-topic) >0.04 not belonging to the locus chr14q32 (see Figure 7).

- A second class of topics is represented by the other three entries of Figure 8 (miRNA-topic 11,13 and 16 in our output) which show a different behavior in one of the subtypes with respect to the others (in the present case these topics are upregulated in samples belonging to the basal subtype). Out of these only miRNA-topic 11 shows a significant entry in the table of enriched keywords Table 2. The enrichment is for another chromosomal locus: chr19q13. What is interesting is that this locus was associated in the past to other types of cancer [55]. Our analysis suggests that it could also play a role in breast cancer and in particular in the Basal subtype. Moreover, we found a not trivial overlap between genes in these miRNA-topics and the miRNA clusters proposed by [56]. In particular there were 12 miRNAs in miRNA-topic 7 from cluster cl349_chr14 (estimating the probability of this happening by chance using an hypergeometric test we obtained a *P* − *value* ≃ 10^−5.8^) and 8 miRNAs in miRNA-topic 11 were assigned with label cl590_chr19 (*P* − *value* ≈ 10^−7.4^).

In the next subsection we shall study in detail, as an example of the type of analyses that we can perform using the probability distributions obtained from triSBM, the first of these topics.

### 2.5. miRNAs contained in miRNA-topic 7 are strongly associated to breast cancer and may affect the survival of patients

We can use the information contained in the probability distribution *P*(miRNA|miRNA-topic) to perform a more refined analysis of the miRNAs contained in the topic. First we see that 75% of the miRNAs in the topic are annotated with chr14q32 locus and that they are exactly those with the highest values of *P*(miRNA|miRNA-topic). This can be visualised in Figure 7 where we highlighted in red the miRNAs annotated to the chr14q32 keyword from GSEA [43].

Then we sorted the miRNAs on the basis of their value of *P*(miRNA|miRNAs’ topic) and investigated the first ones (see those with *P*(miRNA|miRNAs’ topic) >0.030 in Table 3), it turns out, using the DISEASES tool [57], that most of them are in some way associated to breast cancer. Let us highlight that mir-511, mir-31 and mir-34c are highly important in this miRNA-topic, nevertheless they do not belong to the c14q32 locus gene set. What is interesting in our analysis, is that it suggests that these miRNAs, which were studied in the past as separated entities, are most probably working together. A better understanding of this cooperative behaviour could be of great importance to fine tune future therapeutic protocols. As a first step in this direction we took advantage of the probabilistic nature of topic modeling to investigate the survival probability of patients.

**Table 3.** micro-RNAs sorted by their *P*(miRNA|miRNA-topic 7). The most important miRNAs in our candidate miRNA-topic. Most of them are well known in literature. The complete list is available at https://github.com/BioPhys-Turin/keywordTCGA/blob/main/brca/trisbm/trisbm_level_1_keyword-dist.csv.

| microRNA | $P(\text{miRNA}\vert \text{miRNA-topic } 7)$ |
| --- | --- |
| hsa-mir-654 | 0.047 |
| hsa-mir-758 | 0.046 |
| hsa-mir-493 | 0.042 |
| hsa-mir-889 | 0.041 |
| hsa-mir-34c | 0.041 |
| hsa-mir-431 | 0.039 |
| hsa-mir-369 | 0.039 |
| hsa-mir-370 | 0.039 |
| hsa-mir-410 | 0.037 |
| hsa-mir-154 | 0.035 |
| hsa-mir-495 | 0.035 |
| hsa-mir-511 | 0.035 |
| hsa-mir-411 | 0.033 |
| hsa-mir-432 | 0.032 |
| hsa-mir-31 | 0.031 |
| hsa-mir-487b | 0.030 |
| hsa-mir-376c | 0.030 |
| hsa-mir-412 | 0.030 |
| ... | < 0.030 |

In particular since a *P*(topic|sample) can be assigned to each patient (sample), it is possible to create cohorts of patients based on the importance of a given topic in their transcriptome.

We ran a Cox [58] model to verify which is the contribution of our topic to the survival probability of patients.

We report in Figure 9 the Kaplan-Meyer curves that we obtained.

**Figure 9.**
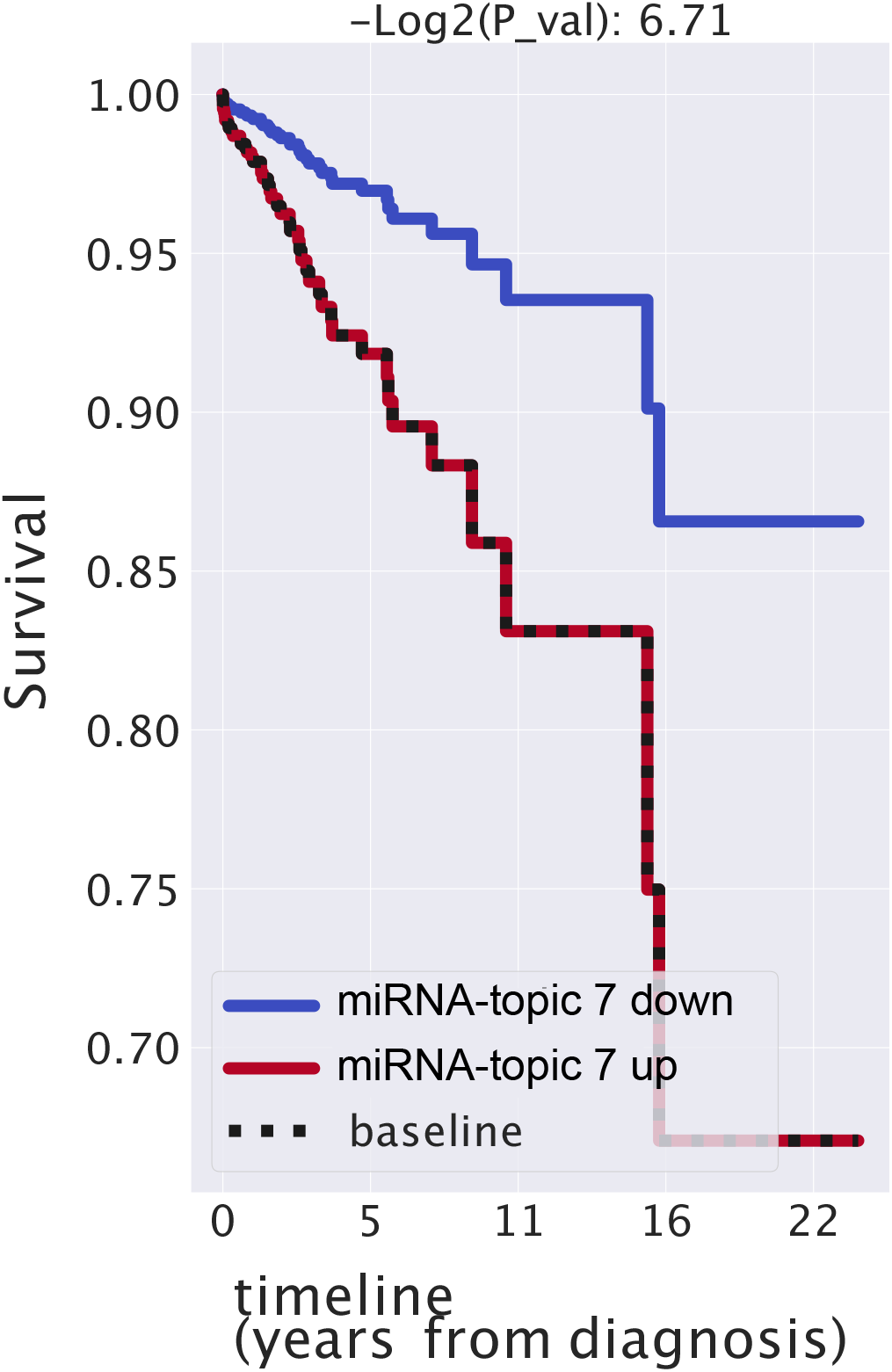
Kaplan-Meier analysis of miRNA-topic 7. We divided patient (samples) into two cohorts using the information the importance of this miRNA-topic in each sample. Patients with a great presence of this topics have smaller values of survival.

The contribution of the topic to the survival probability turns out to be very significant: a positive regulation corresponds to higher hazard ratios meaning that if miRNA inside our topics are expressed higher than normal, the survival probability of patients decreases. While these results should be taken with some caution, due to the several sources of bias which may be present in TCGA population that we tested, it is nevertheless interesting to notice that the presence or absence of this topic has an impact on the survival probability larger than the tumor stage, which is, obviously, strongly correlated with the patient’s prognosis (see Supplementary Figure S3). As a comparison we also report in Figure S3 variables like gender (this is not very balanced, samples are almost all females) or age, which, as expected, do not have a significant effects on the survival probability of patients.

Going further in the investigation of the survival probability of the patient one can wonder if patients in a cluster share a similar prognosis.

If one measures the fraction of patients still alive 3 years after the diagnosis it is possible to give a prognosis indication of patients in a given cluster. In Figure S7 we reported two clusters in which the prognosis of the patient is significant. In cluster 6, for instance, only 18% of the patients survived more than 3 years. This corresponds to a cluster with a bad prognosis. On the opposite side, more than 60% of patients grouped in cluster 14 survived: we can assert that patients in this set have a favourable prognosis. We measured the significance of this results comparing the aforementioned percentages to the ones obtained creating clusters at random (picking up patients from the whole dataset at random 100 times) and obtained significant *Z* ~ 3 scores (reported in Figure S7).

## 3. Discussion

There are two main directions in which the analysis discussed in the previous section could be improved. First, one would like to include in the game the regulatory interactions among miRNAs and target genes. Second, one would like to extend the integration to other information layers. We shall discuss in this section a few preliminary attempts in these directions.

### 3.1 Including regulatory interactions in the triSBM framework

MiRNAs exert their biological function by regulating target genes at the post-transcriptional level. It is thus of great importance to be able to include this information in the topic modeling analysis. This is not an easy task, since miRNAs act in a combinatorial way: typically several miRNAs cooperate to regulate a single target gene and at the same time a single miRNA can regulate hundreds of targets.

Moreover, while the standard miRNA-target regulatory interaction is of inhibitory type, it sometimes happens that a miRNA can have a widespread (indirect) activatory role by interfering with a repressed epigenetic pathway. These are the so called “epi-miRNAs”[59,60] which have been recently shown to play an important role in cancer development [60]. Keeping track of these interactions can be of crucial importance to correctly decode the information contained in the miRNA expression data. To this end one can make use of a few specialized databases of miRNA-target interactions. In particular, in the following we shall use MirDip [61] and TarBase [62], which are among the most popular ones and are somehow complementary in their target selection choices.

To integrate the regulatory information we made use of the analogy of this problem with inclusion of the citations information among documents in standard topic modeling applications to texts [23]. In our case the additional links are not between samples (as it would be a citation link or an hyperlink), but therefore links between branches and in particular we added gene-miRNA links.

We ran the tri-partite model as described before, then, in a second moment, we added links gene-miRNA from regulatory network (we tested separately MirDip [61] and TarBase [62]), as shown in Figure 10a. On the fitted triSBM model we ran steps of the fast merge-split implementation of SBM [63] to improve the description length (see Methods for a precise definition) of the data made by the model taking advantage of the gene-regulation information in a way similar to the citation between documents when they are used to improve the classification ability of hSBM in that context.

**Figure 10.**
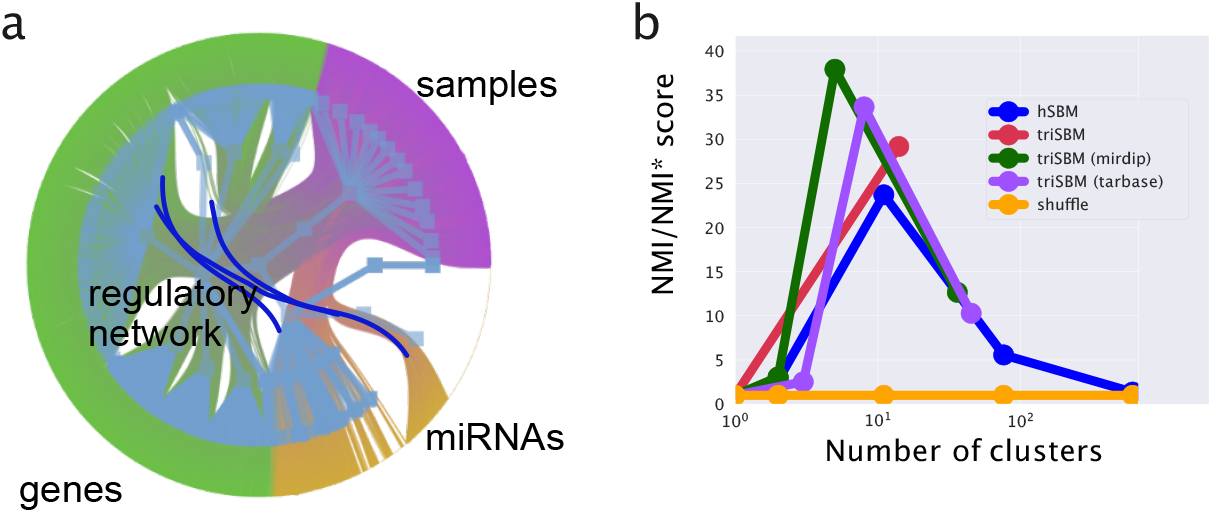
Configuration and scores when adding gene-miRNA links. In (**a**) a cartoon of a tri-partite network with links gene-miRNA. In (**b**) the scores of this new setting using two different (mirDIP [61] and TarBase [62]) regulatory networks separately.

We report in Figure 10b the Normalised Mutual Information measuring the ability of the full process (fit triSBM, add links, run merge-split) in identifying the breast subtypes. Remarkably enough, we see that including the information on miRNA-genes interactions we reach a higher NMI, i.e. a better agreement of our clusters with the subtype organization. This does not happen simply running merge-split after triSBM is run.

This shows that it is possible to integrate not only multiple layers of sample related information, but also knowledge about correlations between different kinds of features. Our results represent a first proof of concept in this direction, and we plan to further pursue this type of analysis in future.

### 3.2. Adding further layers of information: the case of Copy Number Variation

As we discussed in the introduction the nSBM algorithm can be extended in principle to any other layer of information on the samples. A natural candidate is Copy Number Variation (CNV). It is well known that chromosomal aberrations are a hallmark of cancer and that several types of cancer are characterized by a well defined set of chromosomal loci whose deletion or duplication can drive the onset of that particular type of cancer. We already noticed that using the information contained in the miRNA branch we could identify two loci whose alteration was known to be associated to the onset of breast cancer. In TCGA database we also have the information on the CNV values for all samples. We included this information by adding a fourth branch to our algorithm (accordingly we shall call it in the following as “tetraSBM”). As a preliminary test we selected only genes with positive CNV (i.e. genes contained in duplicated loci) and neglected for the moment deletions.

We performed a gene selection also in this new branch. Highly Copied Genes were selected, keeping the ones with an average (over samples) CNV greater than 3.5. 1 353 genes passed our selection. This selection would select genes with at least 2 duplication (CNV = 4), on average.

It is important to stress that at this stage nodes which correspond to the same gene in the gene expression branch and in the CNV branch are completely uncorrelated and are seen by the algorithm as independent nodes. We shall discuss below how to address this issue.

In our setting we have 3 000 protein coding genes in the gene expression branch, 1353 genes in the CNV branch and 417 of them are represented by nodes in both branches.

We ran the tetraSBM model on this network with samples, protein-coding genes, miRNAs and CNV genes and obtained two hierarchical levels. In the first one the four branches were partitioned in 13 clusters, 7 gene-topics, 5 miRNA-topics and 5 CNV-topics. In the second one we found 397 clusters, 49 gene-topics, 14 miRNA-topics and 31 CNV-topics.

Looking at the CNV-topics, we found a very interesting result (see Table 4). Performing the usual Gene Set Enrichment Analysis we found, with very low values of False Discovery Rate (FDR), a few chromosomal loci which, we think, represent the complete collection of chromosomal aberration associated with breast cancer and could be used as a robust signature of this type of tumor. The relevance of this result is supported by the other set of enriched keywords (taken from [64]), which are reported in Table 4 which show that for some of these loci the association with breast cancer was already known and is indeed very strong.

**Table 4.**
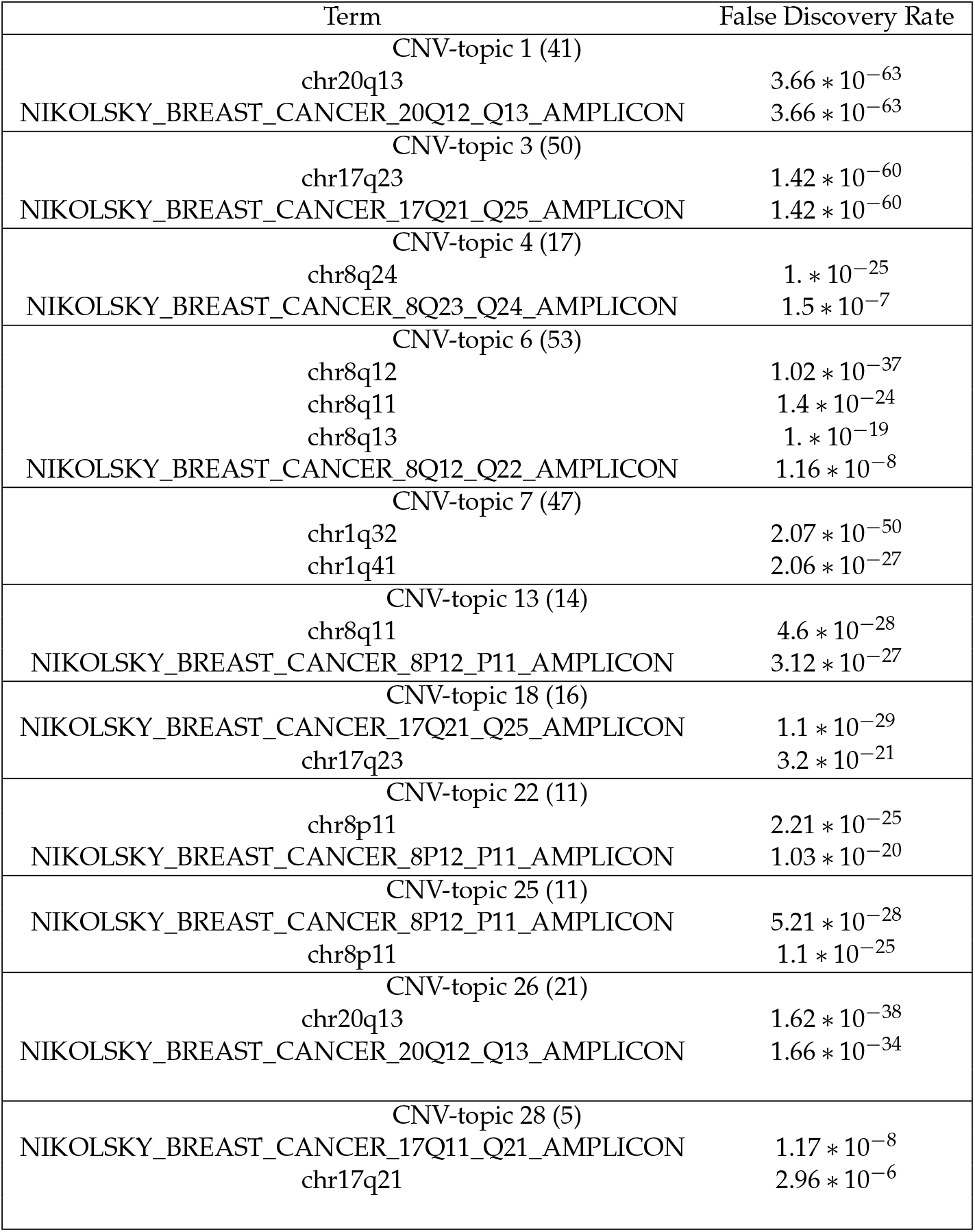
Enrichment analysis on the Copy Number Variation branch of tetraSBM. All the lists are available at https://github.com/BioPhys-Turin/keywordTCGA/blob/main/brca/tetrasbm/trisbm/trisbm_level_0_kind_3_metadata.csv.

| Term | False Discovery Rate |
| --- | --- |
| CNV-topic 1 (41) |  |
| chr20q13 | $3.66 * 10^{-63}$ |
| NIKOLSKY_BREAST_CANCER_20Q12_Q13_AMPLICON | $3.66 * 10^{-63}$ |
| CNV-topic 3 (50) |  |
| chr17q23 | $1.42 * 10^{-60}$ |
| NIKOLSKY_BREAST_CANCER_17Q21_Q25_AMPLICON | $1.42 * 10^{-60}$ |
| CNV-topic 4 (17) |  |
| chr8q24 | $1. * 10^{-25}$ |
| NIKOLSKY_BREAST_CANCER_8Q23_Q24_AMPLICON | $1.5 * 10^{-7}$ |
| CNV-topic 6 (53) |  |
| chr8q12 | $1.02 * 10^{-37}$ |
| chr8q11 | $1.4 * 10^{-24}$ |
| chr8q13 | $1. * 10^{-19}$ |
| NIKOLSKY_BREAST_CANCER_8Q12_Q22_AMPLICON | $1.16 * 10^{-8}$ |
| CNV-topic 7 (47) |  |
| chr1q32 | $2.07 * 10^{-50}$ |
| chr1q41 | $2.06 * 10^{-27}$ |
| CNV-topic 13 (14) |  |
| chr8q11 | $4.6 * 10^{-28}$ |
| NIKOLSKY_BREAST_CANCER_8P12_P11_AMPLICON | $3.12 * 10^{-27}$ |
| CNV-topic 18 (16) |  |
| NIKOLSKY_BREAST_CANCER_17Q21_Q25_AMPLICON | $1.1 * 10^{-29}$ |
| chr17q23 | $3.2 * 10^{-21}$ |
| CNV-topic 22 (11) |  |
| chr8p11 | $2.21 * 10^{-25}$ |
| NIKOLSKY_BREAST_CANCER_8P12_P11_AMPLICON | $1.03 * 10^{-20}$ |
| CNV-topic 25 (11) |  |
| NIKOLSKY_BREAST_CANCER_8P12_P11_AMPLICON | $5.21 * 10^{-28}$ |
| chr8p11 | $1.1 * 10^{-25}$ |
| CNV-topic 26 (21) |  |
| chr20q13 | $1.62 * 10^{-38}$ |
| NIKOLSKY_BREAST_CANCER_20Q12_Q13_AMPLICON | $1.66 * 10^{-34}$ |
| CNV-topic 28 (5) |  |
| NIKOLSKY_BREAST_CANCER_17Q11_Q21_AMPLICON | $1.17 * 10^{-8}$ |
| chr17q21 | $2.96 * 10^{-6}$ |

On the other side, if we test the performance of tetraSBM to identify the samples subtype we see that, including the information on CNV, we have a *decrease* of the NMI value (see the Supplementary Figure S4). This is not surprising because within the duplicated (or deleted) loci, besides the few drivers of the cancer there are hundreds of “hitchhikers” genes which simply add noise to the process of subtype classification performed by the other two layers (genes and miRNAs). The variability of the gene expression values which are associated to the different cancer subtypes (and in fact allowed to classify the subtypes in the hSBM and triSBM versions of the algorithm) were completely shadowed by the noise induced by the CNV branch. In the Supplementary Figure S5 we reported a bi-partite analysis on subtypes with a bi-partite network using only the CNV data. This analysis confirms that the CNV layer is less informative than the layer with only protein coding genes.

This tells us that adding further layers of information does not automatically improves the quality of clustering. It is always important to perform a careful analysis of the biological information contained in the data and of its possible interference with the other layers. In this particular example we learnt that miRNAs cooperate together to assign to the same gene-topic co-regulated genes and in the same clusters samples of the same subtype. This fact becomes particularly clear looking at the probability (see Equation 2 in the Methods section and [65] for further details) of moving nodes between groups: when moving a gene between gene-topics it is more probable to move in a topic where there are genes with many connections to the miRNAs connected to the gene itself. This is confirmed by the fact that, as we discussed in the previous sections, there are miRNA-topics which overlaps with clusters of miRNA [56] known to co-express in breast cancer. On the other hand, the CNV features force samples with the same duplicated loci to be together and this seems not to be correlated with the cancer subtype, at least in TCGA-BRCA data.

This does not mean that the addition of CNV data is useless. It is only by including CNV that we may have, as we have seen, a precise information on the chromosomal aberrations involved in breast cancer. It is also interesting to notice that this information is somehow complementary to the one we obtained in the previous section looking at the miRNA clusters. The chromosomal loci that we detected there, are not present in this CNV analysis, because their CNV value is below the threshold we fixed to include CNVs in the tetraSBM.

## 4. Materials and Methods

### 4.1. The Cancer Genome Atlas data

The results published here are in part based upon data generated by The Cancer Genome Atlas (TCGA) managed by the NCI and NHGRI. Information about TCGA can be found at https://cancergenome.nih.gov. TCGA data of breast cancer samples were downloaded through portal.gdc.cancer.gov. We selected *TCGA* program, *TCGA-BRCA* Project Id, *transcriptome profiling* as Data Category. We chose *Gene Expression Quantification* and *RNA-Seq* as Data Type and Experimental Strategy to download gene expression data in HTSeq-FPKM. Moreover, we downloaded the number of reads per million of miRNA mapped from *miRNA Expression Quantification* Data Type generated with *miRNA-Seq Experimental Strategy*.

#### Metadata and cancer subtypes

In order to benchmark our results we compared in the Results section the clusters of samples obtained by our algorithm with TCGA annotation which we considered as our “ground truth”. We choose the annotations available trough TCGABiolinks [32,33] and in particular the one defined as *Subtype_Selected*. Those subtype annotation are provided by [66] and are based on previously published studies [17,67] about breast cancer based on TCGA.

In other analyses we needed to known if a sample was a primary tumor or derived from normal tissues. Solid Normal Tissues samples are the ones with *sample type* 10 or 11 in TCGA barcode (10 to 19 are normal types) (https://docs.gdc.cancer.gov/Encyclopedia/pages/TCGA_Barcode/).

We downloaded the independent Breast Cancer Consensus Subtypes (BCCS) related to the TCGA files provided by the Supplementary files of [41].

### 4.2. METABRIC miRNA landscape data

We downloaded METABRIC data from European Genome-Phenome archive.

We downloaded METABRIC miRNA landscape study (EGAS00000000122), in particular Normalized miRNA expression data (EGAD00010000438) and Normalised mRNA expression (EGAD00010000434).

### 4.3. nSBM: a multi branch Stochastic Block Modeling algorithm

We collect here some further information on the nSBM algorithm.

- The search of the optimal allocation of the latent variables is performed inheriting and expanding [25] hierarchical Stochastic Block Modeling (hSBM) introduced in [10]. Note that the training process is performed simultaneously in all the branches of the network: this means that all the types of data contribute to the learning process at the same time, without, in principle, any preference at the beginning.
- As mentioned in the main text, nSBM tries to maximise the posterior probability 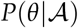 that the model describes the data

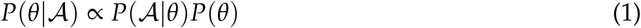

in a completely non-parametric [68] way. Instead of maximizing the probability the model, as usual, it minimises the Description Length 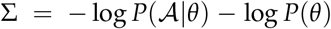. We used the `minimise_nested_blockmodel_dl` function from graph-tool [69]. In our setting 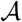 is a block matrix in which each block is a “Bag of Features” (i.e. genes, miRNAs,…). It can be seen as a two dimensional matrix whose entries *w_ij_* are the weights mentioned above. The probability of accepting the move of a node with a neighbour *t* from group *r* to group *s* is [65]

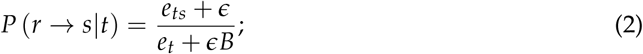

being *e_ts_* the number of edges between groups *t* and *s* and *e_t_* the total number of edges connected to group *t*. From this, it should be clear another advantage of a multi-branch approach: different ‘omics may have their own normalisation. In fact, when moving a sample from *r* to *s* the probability is estimated considering only the branch to which *t* belongs. If the node *t* is a gene *e_ts_*/*e_t_* is normalised taking only into account the mRNA expression values.
- We set the algorithm so as to do a sort of model selection minimising the Description Length Σ 10 times and then choosing the model with the shortest Description Length.
- We used the nested, degree corrected [68], version of the model [70], so as to obtain in output a hierarchy of results.
- The intrinsic complexity of typical Stochastic Block Modeling algorithms is 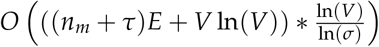 (*τ*, *n_m_* and *σ* are hyper-parameters of the model) which equals *O*(*V* log^2^*V*) if the graph is sparse (*E* ~*O*(*V*)) [71], being *V* the number of vertices (samples, genes and microRNAs) and *E* the number of edges. If *E* >> *V* the complexity is not logarithmic and the CPU time needed to minimize the description length increases as well. In this case, to reduce the CPU bottleneck, one can apply a log-transformation to the data, which strongly reduces the number of edges *E*. We ran the model on a 48-core machine with 768GB of memory [72].

In our setting, we have *V* ~*O*(1000) vertices, and *E* ~ *O*(1000000) edges and the network is indeed very dense, In order to reduce the number of nodes and edges a preprocessing step is needed. We shall discuss this issue in the next subsection.

We considered 1222 samples from TCGA-BRCA project and selected the 1200 with a valid annotation from [32,33]; then we ran the model on a tri-partite network built with normal and tumor samples from TCGA on one branch, 3 000 FPKM normalised gene expression data on a second branch and 1300 miRNA-Sequencing data on the third branch. Note that we did not explicitly selected the known breast Cancer markers, our approach to topic model, as already discussed in [3], took into account the whole expression pattern and does not relay only on few specific markers.The output of the tri-partite model consisted of two hierarchical levels with 1 and 14 clusters; 11 and 331 topics; 33 and 47 miRNA-topics on the three branches respectively. We ran also, as a comparison without miRNAs, hSBM on a bipartite network and we obtained levels with 2,11, 76, 608 clusters and 5,17, 62, 390 topics across the hierarchy.

As output of the model, we find the probability distributions *P*(topic|sample) and *P*(gene|topic). These probabilities are defined, in terms of entries of the program as follows:

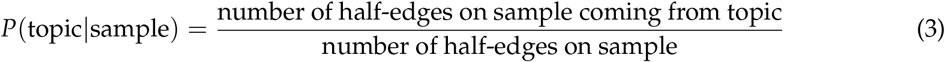

and

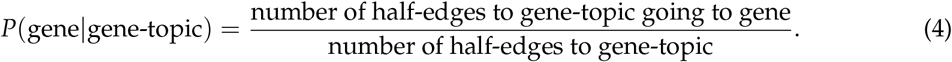

(The same is true for miRNA-topics and for each end every eventual additional layer of features.

#### Gene and miRNA selection

The data provided in the atlas consisted of 1222 (~ 1100 have both mRNA and miRNA transcript profiles data) samples associated to almost 20 000 genes and 2 000 miRNA entries. Without preprocessing this would have led to an adjacency matrix too big to be handled efficiently by the algorithms.

We performed two kind of preprocessing to reduce the number of nodes and the number of edges.

In order to reduce the number of nodes we filtered genes and miRNAs selecting only the highly variable ones. The highly variable are the ones with the highest dispersion (variance over mean) with respect to the genes with the same average expression. This selection was performed using *scanpy* python package [73]. This analysis was performed separately on genes and microRNAs since they are provided by different experiments and different normalisation. We selected in this way 3 000 genes and ~ 1 200 miRNAs.

Furthermore, we applied a standard approach to reduce the weights of the links and applied a log(*FPKM* + 1) transformation to the data before running the topic models. This helped us to reduce by some order of magnitudes the number of edges (as we mentioned above in this class of algorithms the weight of a link is mimed adding multiple edges with weight 1) and the model ran times faster.

In the Copy Number Variation analyses we choose ~ 1300 genes with an average CNV >3.5.

An interesting feature of SBM type of algorithm is that they are typically robust with respect to gene selection. In the analyses of this paper we considered only highly variable genes but in the supplementary material of [3] we discussed different types of gene selections showing that they were typically leading to similar performances.

In the analysis of METABRIC dataset we utilised the previously selected genes and microRNA.

#### Evaluation metrics

To evaluate the agreement between the sample partitions and the annotations we choose the so called “Normalised Mutual Information” (NMI) which was proposed in [40] in a new evaluation framework for topic models. Moreover, as discussed in [3] it can be shown that NMI is the harmonic average of two metrics which evaluate respectively the completeness and the homogeneity of a partition of annotated samples [39]. A cluster is complete if all samples with a given label are assigned to the same cluster; a partition is homogeneous if in a cluster all the samples have the same annotation. In order to correctly identify the cancer subtype of a given sample one would like to have a partition in clusters which is both homogeneous and complete.

The *NMI* is estimated using the Shannon’s entropy formula to measure the quantity of information in the partition. The problem of this measure is that even in a random partition there is a residual entropy and the *NMI* is not zero, this effect is particularly important in the layers of the models with high resolution (many clusters). In order to avoid this bias, we evaluated this default *NMI* by randomizing the subtype annotations of the samples. This was done multiple (~50) times preserving each time the number of clusters and the number of samples in every cluster; we call the average *NMI* on this multiple random assignments *NMI*\*, this is the residual information on the considered partition. In the results we reported NMI/NMI* which measures how much information the model learns with respect to a random assignment. It is important to stress that this measure has not an absolute value and should not be used to compare performances on different datasets, however it can be successfully used to compare different algorithm in the same dataset, which is what we did in the Results section.

##### Description length Σ how well the model is describing the data

In addition to the NMI it is also possible to compare different classes of topic modeling algorithms on their ability to compress the data [65,74]. This can be addressed measuring the description length Σ of a model, which represents, in nat units, the number of bits a model requires to describe the data network. Unlike NMI, it has the advantage not to rely on any ground truth. Using 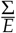 (being *E* the total number of edges) it is possible to measure the quantity of information taht the model requires to describe an edge. In the models of Figure 5 hSBM requires a 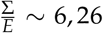 which is greater than the 1,4 units required by triSBM. One can estimate the difference of the two 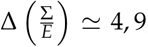, this can be related to the Bayes factor [75] (being the posterior *P* = exp – Σ) Λ = exp ΔΣ ≃ *e*^4,9^ ≃ 10^2,1^ it means that the model with miRNA is a ~ 100 time more probable description of the data network links. The description lengths of the tetra-partite model and the model with regulatory network are reported in Supplementary Materials (see Figure S6).

### 4.4. Construction of the 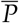(topic|subtype) distributions

From the *P*(topic|sample) distributions it is easy to obtain the probability *P*(topic|subtype) by averaging over all samples belonging to the same subtype.

Then, by subtracting to *P*(topic|subtype) the mean value over the whole dataset we find a new set of quantities, that we define as “centered” distributions (we already used them in [3] and it has the same meaning of the normalised value of the mixture proportion *τ* in [23])

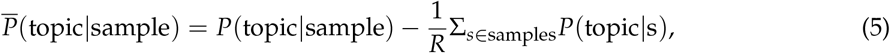

being *R* the total number of samples. This procedure can be implemented separately both on the miRNA-topic and on the gene-topic side. The centered *P*(topic|sample) can be represented as box plots, after grouping samples by their subtype. Examples of these are the box plots reported in Figure 6 on the gene side and Figure 8 on the miRNA side.

#### Survival analysis

We performed the survival analyses fitting a COX [58] model.

Our analysis began with the list of the mixtures *P*(topic|sample). We cleaned up the stages’ labels removing any additional letter (e.g., *stage ia* became *stage i*) and ended up with four stages *i, ii, iii* and *iv*.

Using Genomic Data Commons tools we downloaded TCGA metadata and in particular: *demographic.vital_status, demographic.days_to_last_follow_up, demographic.days_to_death, demographic.gender* and *diagnoses.age_at_diagnosis*. We estimated the lifetime or the number of days the patient survived after the diagnosis, using *days_to_last_follow_up* if the patient was *Alive* and *days_to_death* for *Dead* patients. A similar approach was recently utilized by [76].

In order to estimate whether a topic is up regulated in a patient, we evaluated the 35th percentile of *P*(sample|topic) and considered it as a threshold *thr*. Then we engineered a feature as follows:

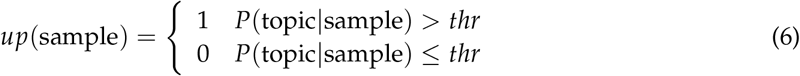

We used these data to fit the hazard with a *COX* model. These analyses were performed using *lifelines* Python package [77], and in particular the COXPHFitter module. We used the lifetime, the vital status and the new feature as input for the fit function.

The Cox model quantified how the topic of miRNAs regulation affected the survival probability. Cox fits the hazard function conditioned to a variable 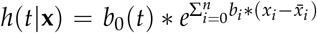. **x** is the vector of the *n* covariates considered. The hazard is defined as the ratio of the derivative of the survival and the survival itself 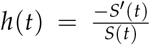. *S*(*t*) is the probability of being alive at time t, namely the number of patient alive at time *t* divided by the total number of patients. The package estimated the ratio between the hazard of samples with topic up-regulated and hazard of samples with topic not up-regulated. Therefore, we were able to estimate the exp(*coef*) or hazard ratio 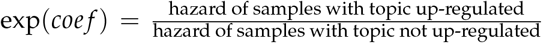. Note that the *coef* does not depend on time, but it is a weighted average of period-specific hazard ratios.

#### Code and nSBM software package

Notebooks to reproduce the results in this work are available on GitHub at https://github.com/BioPhys-Turin/keywordTCGA

The Python package to run nSBM [25] can be downloaded from GitHub (https://github.com/BioPhys-Turin/nsbm) or, alternatively, it can be installed using Anaconda (https://anaconda.org/conda-forge/nsbm) running `conda install nsbm -c conda-forge`.

We discussed in the paper the application using genomics data, however the package is written in a way that makes it agnostic with respect to the kind of data it receives in input and to the number of branches. One can ideally integrate as many different sources (‘omics) of data as needed. Eventually it can process not only biological data, but every kind of dataset whose input could be represented as a rectangular matrix (Bag of Words) for each feature.

## Conclusions

In conclusion the nSBM model we propose here, integrating multiple sources of information into an hSBM analysis, should be useful to extract a lot of information from transcriptomics data. In particular:

- using the python package: *nSBM*, inherited from hSBM [10], ready to install and easily executable on n-partite networks, will be straight-forward to address different types of biological data.
- Second, the integration of multiple sources of data such as microRNA expression levels and the protein coding mRNA ones, greatly improves the ability of the algorithm to identify breast cancer subtypes.
- Third, we use our results to identify a few genes and miRNAs and characterize a few chromosomal duplications which seem to have a particular prognostic role in breast cancer and could be used as signatures to predict the particular breast cancer subtypes.

In conclusion, this paper released a new tool to easily integrate different sources of data into a topic-modeling analysis.

We showed some application in a specific case (breast cancer) with some sources of data (mRNA, miRNA, CNV). Indeed this approach can be applied to other datasets and more important to any possible sources of data (genomics, proteomics, lncRNA, circRNA…).

## Supplementary Materials

The following are available online at http://www.mdpi.com//1/1/0/s1,

Figure S1. Normalised Mutual information of hSBM and triSBM partition compared to the Breast Cancer Consensus Subtypes of Ref. [41].

Figure S2. Validation on METABRIC dataset.

Figure S3. Multivariate (Log) Hazard Ratios.

Figure S4. Normalised Mutual Information of models with samples and mRNA (hSBM), plus miRNA (triSBM) and mRNA plus both miRNA and CNV (tetraSBM). Adding CNV introduces noise to the model.

Figure S5. Normalised Mutual Information of bi-partite models with samples and mRNA (hSBM) and samples with Copy Number Variation (CNV). Adding CNV introduces noise to the model.

Figure S6. Description length of different settings.

Figure S7.Days of survival of different patients in clusters.

## Author Contributions

conceptualization, F.V., M.O. and M.C.; methodology, F.V., M.O. and M.C.; software, F.V.; writing—original draft preparation, F.V., M.C.; writing—review and editing, F.V., M.O. and M.C.; visualization, F.V. All authors have read and agreed to the published version of the manuscript.

## Acknowledgments

We would like to acknowledge the Competence Centre for Scientific Computing C^3^S which provided us the access to the computing cluster OCCAM. The results shown here are in part based upon data generated by TCGA Research Network: https://www.cancer.gov/tcga.

## Abbreviations

The following abbreviations are used in this manuscript:

SBM: Stochastic Block Modeling
TCGA: The Cancer Genome Atlas
GSEA: Gene Set Enrichment Analysis
FDR: False Discovery Rate
FPKM: Fragments Per Kilobase of transcript per Million mapped reads

