## Supplementary Information for "Multi-omics Topic Modeling for Breast Cancer Classification"

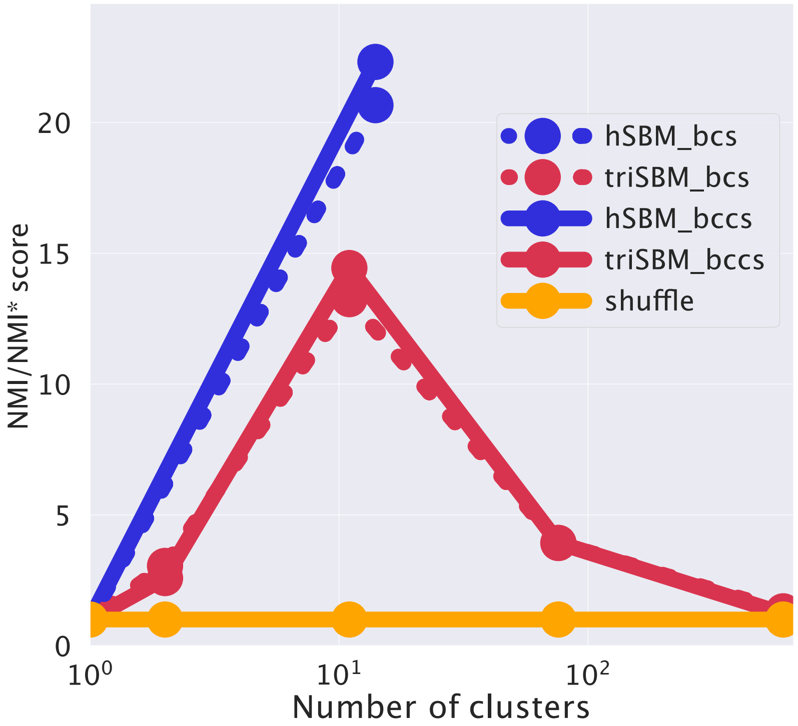


Figure S1: **Normalised Mutual information of hSBM and triSBM partition compared to the Breast Cancer Consensus Subtypes of Horr et al..** Dashed lines correspond to the BCS labels, continuous lines are BCCS labels. The blue lines correspond to triSBM model which integrates the miRNA branch, red lines are hSBM model on bi-partite network. In the models which integrate miRNA into the analyses there is a better agreement between our clusters and the labels provided by the aforementioned independent study.

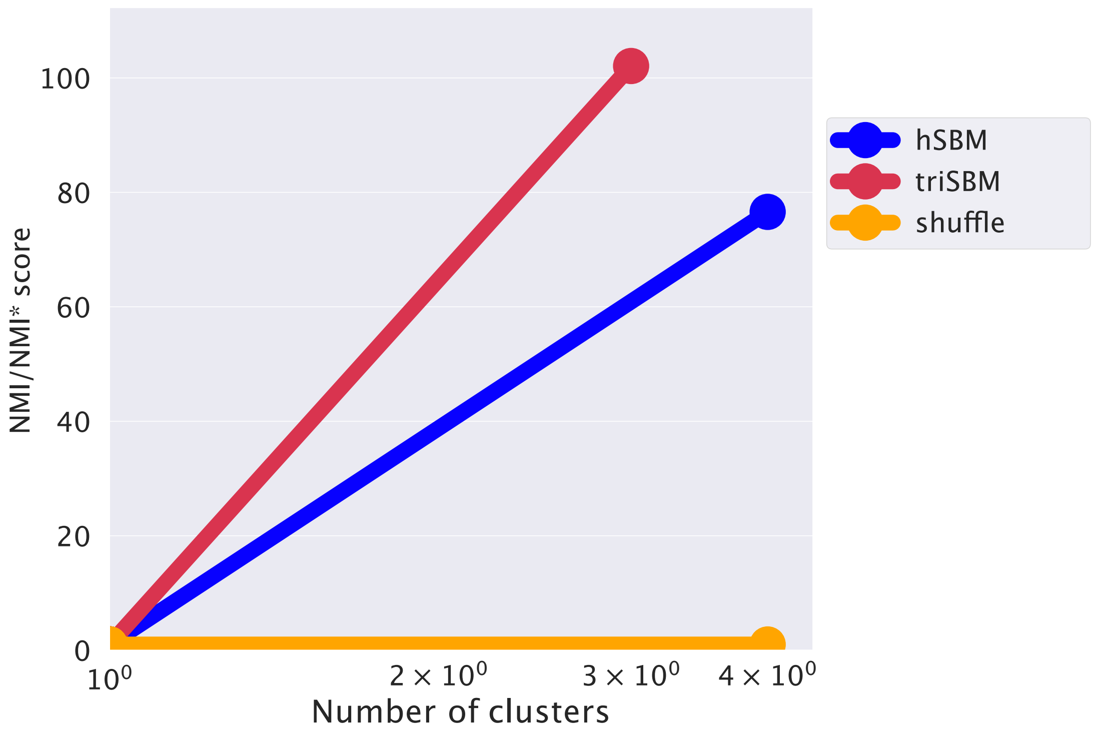


Figure S2: **Validation on METABRIC dataset.** We run hSBM bi-partite model and triSBM model including both mRNA and miRNA data on METABRIC dataset. As reported here the improvement given by the introduction of an additional feature can be seen also on this dataset.

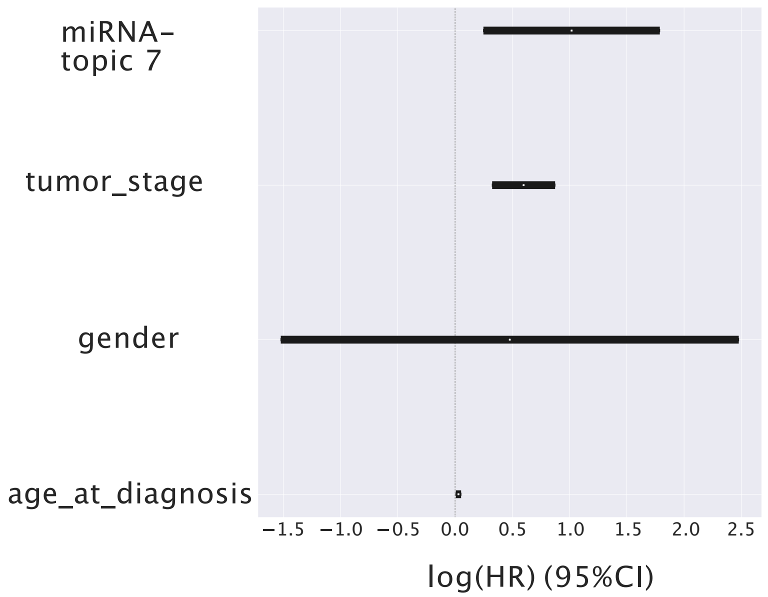


Figure S3: **Multivariate (Log) Hazard Ratios.** We report our candidate topic compared to other covariates whose contribution may correlates with the hazard ratio.

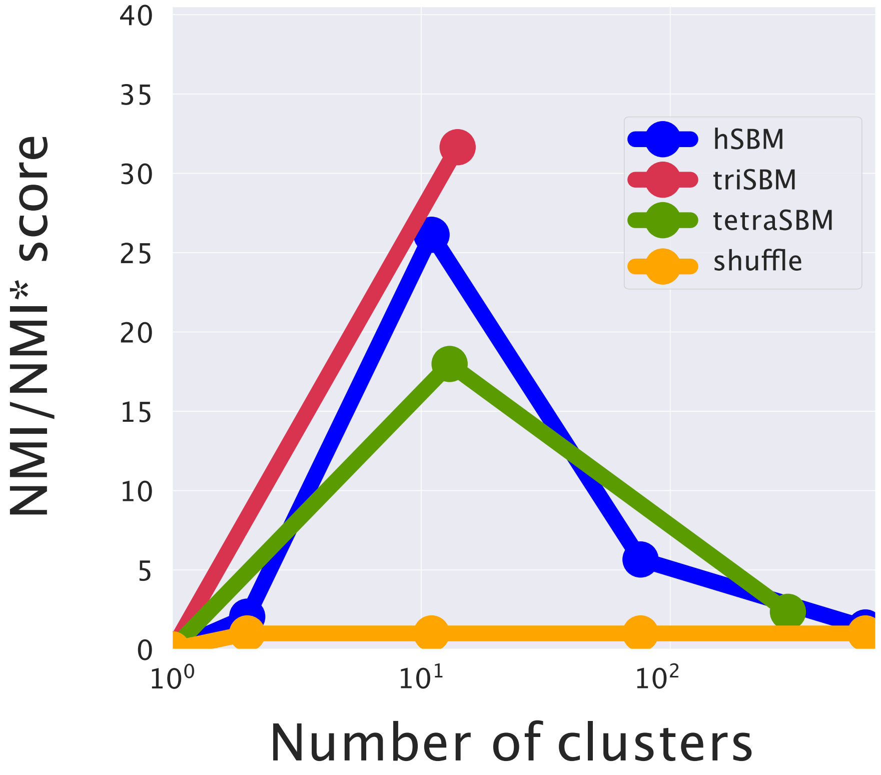


Figure S4: **Normalised Mutual Information of models with genes and mRNA (hSBM), plus miRNA (triSBM) and plus both miRNA and CNV (tetraSBM).** Adding CNV introduces noise to the model.

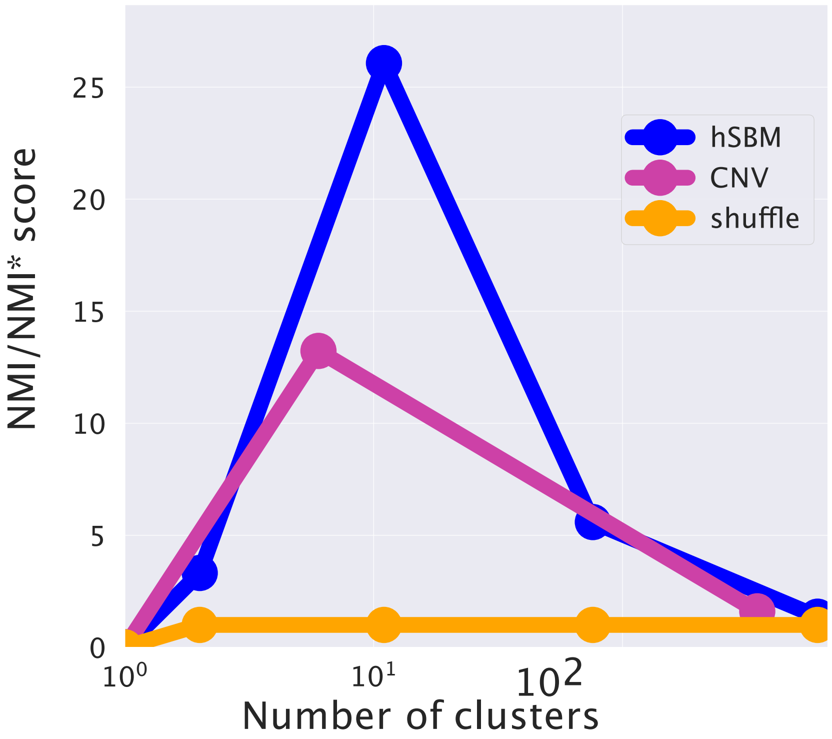


Figure S5: **Normalised Mutual Information of bi-partite models with samples and mRNA (hSBM) or samples and Copy Number Variation (CNV).** Adding CNV introduces noise to the model.

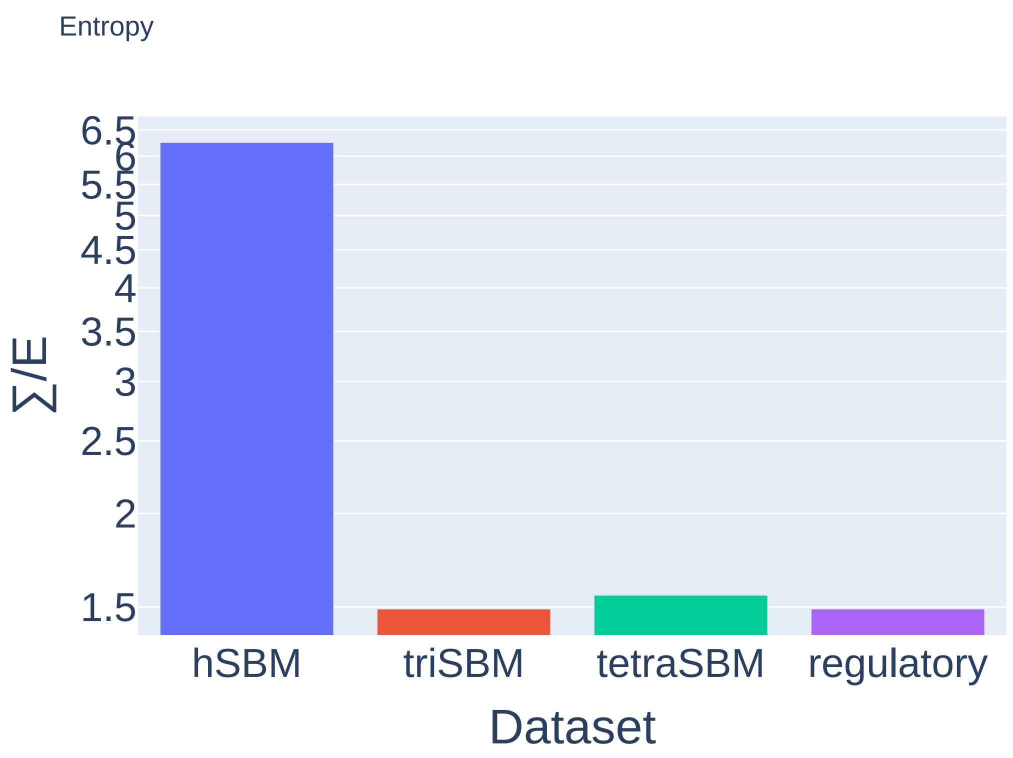


Figure S6: **Description length of different settings.** The probability the data $X$ are described by a model $\theta$ can be written as $P = \exp -\Sigma$, being $\Sigma$ the so-called description length, the lower the better the model describes the data. Here we report description length per edge $\frac{\Sigma}{E}$ in different settings.

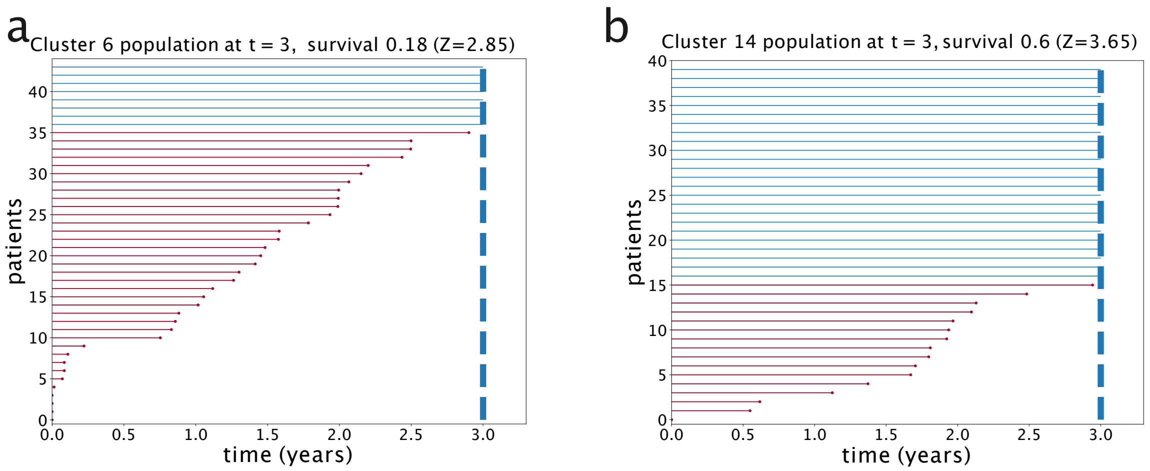


Figure S7: **Days of survival of different patients in clusters.** Each plot represent a cluster, each line corresponds to a patient; the length of the line is proportional to the number of days the patient survived after diagnosis. In one cluster there may be mostly patients died before 3 years (a) or ones who survived after 3 years (b). This result is significantly different from a result obtained shuffling those particular clusters the patients at random (see $Z$ in the Figure).
